# Metabolic engineering of *Saccharomyces cerevisiae* for improved polyphosphate production

**DOI:** 10.64898/2026.11.03.763959

**Authors:** Makarius Baier, Tobias Karmainski, Eva Miriam Buhl, Alexander Deitert, Roy Eerlings, Frederik Völker, Lennart Brans, Evelyn Deis, Philipp Demling, Lars M. Blank

## Abstract

Finite high-grade phosphate rock reserves and rising legislative demand for phosphorus recycling create a need for processes converting phosphorus-containing residual streams into valuable products. Microbial synthesis of inorganic polyphosphate (polyP) could complement its energy-intensive chemical manufacturing while enabling phosphorus recovery in a value-added product with applications ranging from food processing to flame retardants and fertilizer. After orthophosphate (P_i_) starvation and refeeding, *Saccharomyces cerevisiae* naturally accumulates polyP as a cellular P_i_ reserve, but the current multistage process complicates industrial implementation. Here, we identified metabolic interventions that uncouple polyP accumulation from P_i_ starvation by systematic evaluation of 57 engineered *S. cerevisiae* CEN.PK113-7D strains. Constitutive activation of the PHO regulon produced the strongest increase in polyP, whereas disruption of inositol pyrophosphate synthesis, P_i_ import, or the vacuolar transporter chaperone complex impaired accumulation. This pattern indicates that efficient polyP accumulation depends on coordinated activation of P_i_ acquisition and vacuolar polymerization. Notably, deletion of *PHO80* converted a transient starvation response into a constitutive production phenotype, enabling high polyP accumulation during growth. Optimization of the growth medium increased the polyP content further, and efficient use of the supplied P_i_ was demonstrated, while electron microscopy paired with energy-dispersive X-ray spectroscopy supported increased vacuolar storage. During aerobic exponential fed-batch cultivation, *pho80Δ* simultaneously formed biomass and polyP, reaching a polyP content of 22.1% of cell dry weight and a titer of 8.2 g/L, both expressed as KPO_3_ equivalents. These findings establish *S. cerevisiae pho80Δ* as a promising platform for biological polyP production and phosphorus recovery.

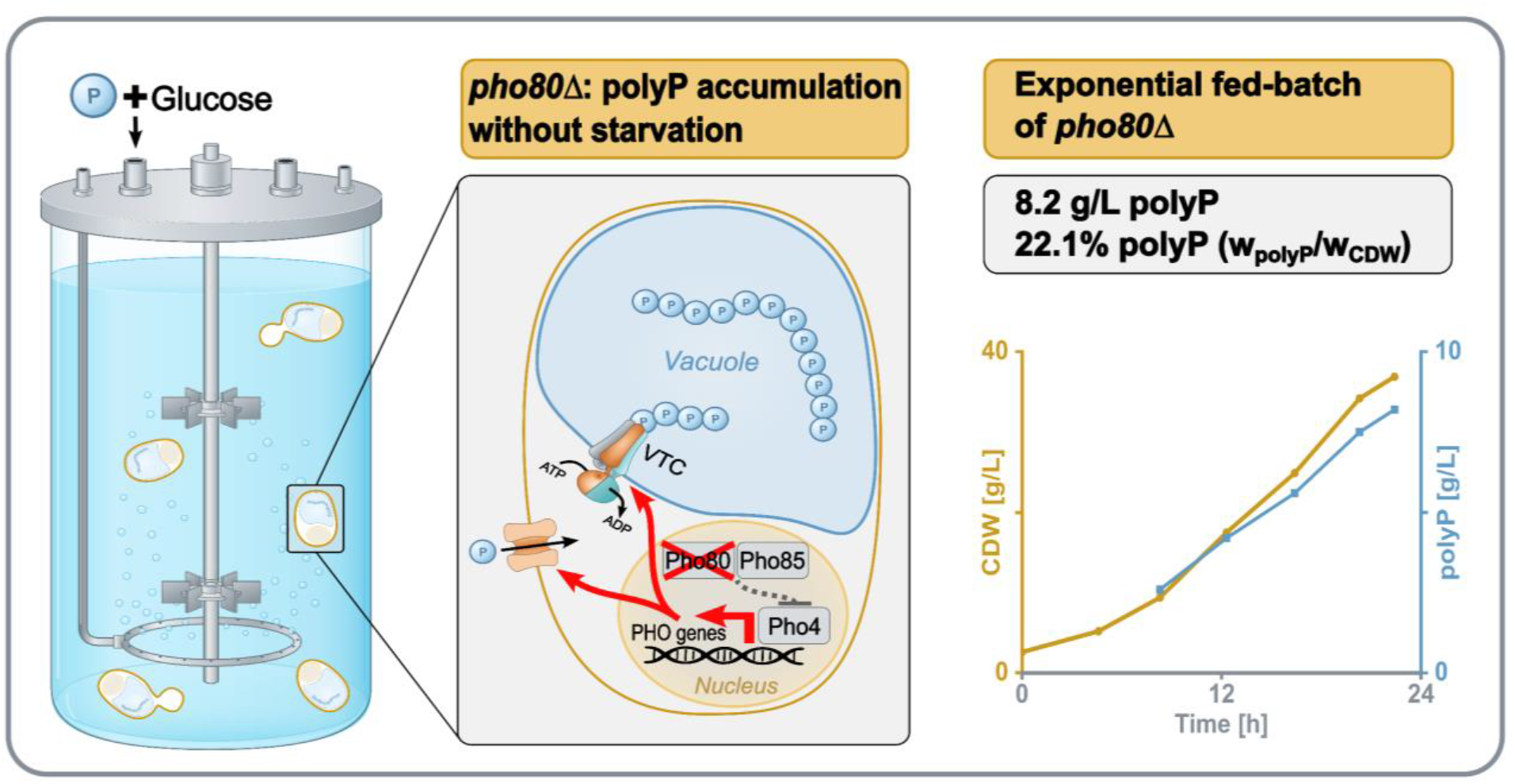

## 1. Introduction

### 1.1 Biological and industrial relevance of orthophosphate (P_i_) and polyphosphate

Phosphorus in the form of P_i_ plays a part in numerous cellular processes and is an essential building block for life as we know it. In the form of fertilizer, humankind has utilized this dependency for the industrial intensification of agriculture. Taken together with the plethora of industrial use cases for phosphorus, this has led to an overuse of this element, raising concerns about depletion of high-grade phosphate rock (Amar et al., 2022) and ecological consequences of eutrophication (Smith and Schindler, 2009). Human alteration of the phosphorus cycle is included within the planetary boundary for biogeochemical flows, one of nine boundaries defining a safe operating space for humanity (Richardson et al., 2023). These concerns have prompted legislative measures to promote phosphorus recovery, including Germany’s revised Sewage Sludge Ordinance (Sichler et al., 2022).

We propose recovering P_i_ from residual streams as value-added, biologically produced inorganic polyphosphate (polyP) as part of a broader effort to support more sustainable phosphorus use.

Among the chemical forms in which living cells store and mobilize phosphorus, polyP is remarkable for both its structural simplicity and its functional diversity. PolyP is a linear polymer composed of P_i_ residues linked by energy-rich phosphoanhydride bonds, with chain lengths ranging from only three to several thousand P_i_ units (Brown and Kornberg, 2008). It is found throughout all domains of life and contributes to numerous cellular processes, including P_i_ and energy storage, stress resistance, and membrane transport (Kulaev, 2004). In vertebrates and humans, polyP additionally participates in specialized processes such as biomineralization and blood coagulation (Kulakovskaya et al., 2012; Kullik et al., 2024).

PolyP is not only a conserved component of cellular P_i_ metabolism but also a chemically versatile polymer with substantial technological potential. Many physicochemical properties of polyP depend on its chain length, allowing different fractions to be used for distinct industrial purposes (Kulakovskaya et al., 2012; Rashchi and Finch, 2000). PolyP provides effective pH buffering, improves water retention in meat products (Shen and Swartz, 2010), and has strong cation-chelating as well as bacteriostatic properties, which are valuable in food preservation and other processes requiring control of metal availability or microbial growth (Lorencová et al., 2012; Younes et al., 2019). PolyP is biodegradable, non-flammable, and generally of low toxicity, supporting its use in products including fertilizers, flame retardants, and food additives (Demling et al., 2024; Kulakovskaya et al., 2012).

Industrial polyP is currently produced predominantly through the thermal condensation of highly purified phosphoric acid obtained from mined phosphate rock (Van Wazer, 1958). This process requires high temperatures and consequently entails substantial energy demand, while also relying on a non-renewable fossil phosphorus resource. Furthermore, common commercial sodium and potassium polyPs typically contain 2–40 P_i_ units, thereby restricting the range of chain-length distributions readily available from conventional production processes (Rashchi and Finch, 2000). Biological polyP production could provide a complementary route operating under milder conditions, with lower energy requirements, the potential to recover phosphorus from diverse P_i_-containing residual streams.

### 1.2 *Saccharomyces cerevisiae* as a platform for biological polyP production

The capacity of microorganisms to take up excess P_i_ from their environment and store it intracellularly as polyP has been recognized for several decades, including early observations in algae and yeasts (Liss and Langen, 1962; Plouviez et al., 2021). Most subsequent work has focused on the genetics, regulation, and physiology of microbial polyP accumulation rather than its exploitation in industrial production processes (Austin and Mayer, 2020; Gerasimaitė et al., 2014). Consequently, few studies have investigated the targeted production of polyP-rich biomass or the recovery of purified, biologically synthesized polyP (Christ and Blank, 2019; Christ et al., 2020).

Those studies identified the yeast *Saccharomyces cerevisiae* as a particularly promising host for biological polyP production. Decades of research on yeast have established large-scale production processes and required infrastructure, and yeast has a long history of safe use in food production, with food-grade strains used in applications generally recognized as safe (Parapouli et al., 2020; Tao et al., 2022). For polyP production in yeast, Christ and Blank found that after P_i_ starvation, subsequent P_i_ feeding can induce rapid P_i_ uptake and strong intracellular polyP accumulation, reaching up to 28% (w/w) of cell dry weight (Christ and Blank, 2019). This phenomenon has variously been described as hyperaccumulation, overcompensation, surplus uptake, luxury uptake, or P_i_ overplus. In this work, the term P_i_ overplus response is used specifically for polyP accumulation induced by P_i_ starvation followed by P_i_ feeding according to Plouviez et al. (2025), whereas P_i_-replete cultivation refers to biomass production under sufficient P_i_ availability. Yeast-based polyP accumulation has also been demonstrated using alternative P_i_ sources, including industrial waste streams and plant-derived residues, illustrating the potential robustness and flexibility of the process (Fees et al., 2022; Herrmann et al., 2023). Efficient exploitation and engineering of these accumulation regimes require an understanding of how *S. cerevisiae* senses intracellular P_i_ availability and coordinates P_i_ acquisition, transcriptional adaptation, and vacuolar storage.

### 1.3 Inositol pyrophosphate (IPP)-dependent coordination of P_i_ acquisition and storage

#### 1.3.1 IPP metabolism and P_i_ sensing

P_i_ is an essential constituent of nucleic acids, phospholipids, and nucleotides, but excessive free cytosolic P_i_ can reduce the free energy available from P_i_-releasing reactions such as ATP hydrolysis. Therefore, cells have to balance P_i_ demand, P_i_ availability, and the concentration of free P_i_. This is achieved by a complex regulatory network that senses P_i_ concentration, governs uptake, storage, and consumption, integrating these with overall metabolism and cell proliferation (Fig. 1).

**Fig. 1:**
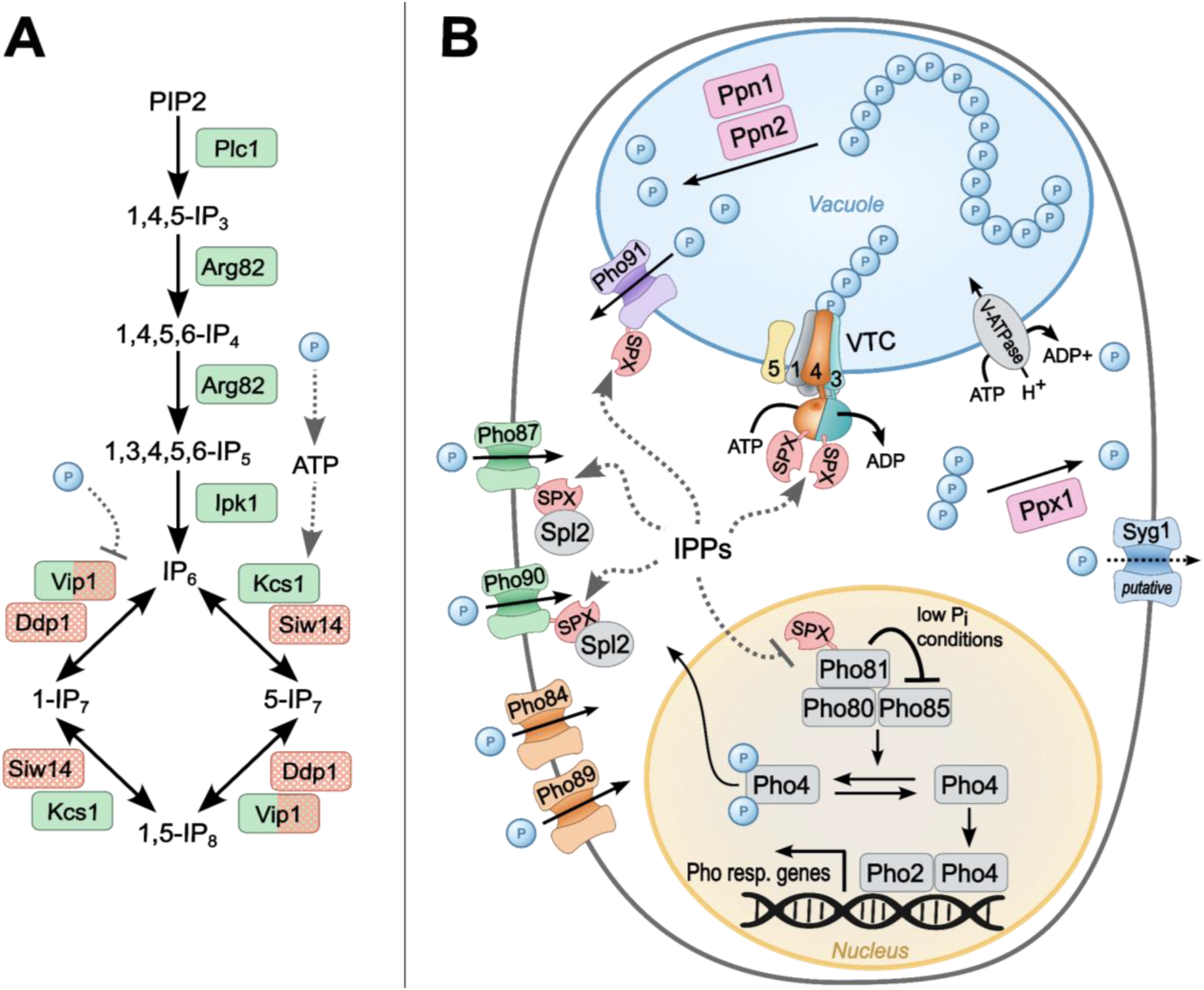
Overview of inositol pyrophosphate signaling and phosphate metabolism in S. cerevisiae. (**A**) Synthesis and interconversion of inositol phosphates (IPs) and inositol pyrophosphates (IPPs). Kinase activity in green; Phosphatase activity in orange (Chabert et al., 2023; Kim et al., 2023; Raia et al., 2025). (**B**) Simplified model of orthophosphate (P_i_) uptake, PHO regulon activation, vacuolar polyphosphate (polyP) synthesis, and polyP degradation. Plasma-membrane transporters supply cytosolic P_i_, while the VTC complex synthesizes polyP from ATP and directly translocates it into the vacuolar lumen (only version containing Vtc3, and not Vtc2 shown). Vacuolar polyP is degraded by Ppn1 and Ppn2, whereas Ppx1 acts on the putative cytosolic polyP pool depicted here. IPPs regulate several SPX-domain-containing components of P_i_ homeostasis, including VTC, Pho81 and P_i_ transporters. Under low-P_i_ conditions, Pho81 inhibits the Pho80–Pho85 cyclin-dependent kinase complex, promoting accumulation of dephosphorylated Pho4 in the nucleus and Pho4–Pho2-dependent transcription of PHO-responsive genes (Austin and Mayer, 2020). Pho4 nuclear-export machinery and transporter-coupled ion transport are omitted for clarity. Spl2-mediated transporter regulation is shown in a simplified form. Syg1 is depicted as a putative P_i_ exporter. Proteins included in the deletion screen but not depicted in the model were omitted to maintain focus on the core regulatory network, as they have indirect or supportive functions, participate primarily in P_i_ scavenging, were selected based on links to other biotechnologically or physiologically relevant pathways, or currently lack a defined mechanistic role in P_i_ homeostasis. Individual blue spheres labeled P represent P_i_, whereas connected spheres represent polyP. Solid arrows indicate transport or metabolic conversion, dotted arrows indicate regulatory or proposed interactions, and blunt-ended lines indicate inhibition.

Inositol phosphates (IPs) and IPPs form a metabolic signaling network to sense P_i_ availability. In the canonical pathway, phospholipase C (Plc1) generates IP₃ from phosphatidylinositol-4,5-bisphosphate (PIP_2_), followed by sequential phosphorylation through Arg82 and Ipk1 to produce IP₆. Kcs1 and Vip1 then jointly synthesize 1,5-IP₈ through two interconnected routes (Chabert et al., 2023).

The abundance of 1-IP₇, 5-IP₇ and 1,5-IP₈ is further controlled by several phosphatases, including Oca5 (Qin et al., 2023), Siw14, Ddp1, and the intrinsic phosphatase domain of Vip1 (Chabert et al., 2023). Direct inhibition of the Vip1 phosphatase domain by P_i_ provides one potential mechanism through which intracellular P_i_ availability could influence the 1,5-IP₈ pool (Raia et al., 2025). In parallel, the ATP dependence of Kcs1 and Vip1 may couple IPP synthesis to cellular energy status, although the quantitative importance of this mechanism *in vivo* remains unresolved (Kim et al., 2023). During P_i_ scarcity, the 1,5-IP₈ pool declines rapidly, potentially reflecting both reduced ATP-dependent kinase flux and increased Vip1 phosphatase activity. Functionally, 1,5-IP₈ regulates P_i_ homeostasis through interactions with SPX-domain proteins, including inhibition of Pho81 and activation of the vacuolar transporter chaperone (VTC) complex, thereby coordinating PHO regulon activity with vacuolar polyP synthesis (Chabert et al., 2023; Pipercevic et al., 2023).

#### 1.3.2 Regulation of the PHO transcriptional response

Under P_i_-replete conditions, 1,5-IP₈ binds to the N-terminal SPX domain of the cyclin-dependent kinase inhibitor Pho81 and functionally inactivates it by destabilizing regulatory contacts between the SPX domain and the Pho80–Pho85 complex, thereby permitting kinase activity. As cytosolic P_i_ becomes limiting, the cellular concentration of 1,5-IP₈ declines rapidly, and the ligand-free SPX domain contributes to activation of Pho81. Under P_i_-replete conditions, active Pho80–Pho85 phosphorylates the transcription factor Pho4 at multiple sites, repressing the PHO regulon through two complementary mechanisms: phosphorylation inhibits nuclear import of Pho4 and promotes its nuclear export, resulting in cytosolic accumulation, while phosphorylation at another site impairs the interaction of Pho4 with Pho2 (Jeffery et al., 2001; Komeili and O’Shea, 1999). Inhibition of Pho80– Pho85 during P_i_ scarcity consequently allows Pho4 dephosphorylation, favors its nuclear import, and reduces its export, leading to accumulation of transcriptionally active Pho4 in the nucleus (Austin and Mayer, 2020). Nuclear Pho4 binds conserved promoter elements and cooperates with the homeodomain transcription factor Pho2, which increases promoter-binding affinity and facilitates recruitment of the general transcriptional machinery. The Pho4–Pho2 complex induces transcription of genes involved in P_i_ acquisition, scavenging, intracellular recycling, and storage, including *PHO5*, *PHO8*, *PHO84*, *PHO89*, *SPL2,* and components of the VTC complex (Austin and Mayer, 2020; Ogawa et al., 2000). Because *PHO81* is itself a Pho4-responsive gene, its induction reinforces the inhibition of Pho80–Pho85 and establishes a positive feedback loop that stabilizes PHO regulon activation during sustained P_i_ limitation (Chabert et al., 2023; Ogawa et al., 2000). Whereas moderate P_i_ scarcity may initiate PHO signaling through an ATP-dependent decline in IPP synthesis, complete P_i_ starvation causes 5-aminoimidazole-4-carboxamide ribonucleotide (AICAR) accumulation as purine biosynthesis stalls, which has been proposed to further suppress IP₈ formation and stabilize the Pho4–Pho2 interaction, thereby reinforcing full induction of the PHO regulon (Kim et al., 2023).

#### 1.3.3 IPP-dependent activation of VTC-mediated polyP synthesis

In parallel with the transcriptional control of the PHO regulon, IPPs directly regulate P_i_ storage through the VTC complex. In *S. cerevisiae*, VTC forms two alternative heteropentameric complexes (3x Vtc1, 1xVtc4, 1x Vtc2 or Vtc3), while Vtc5 can associate as an accessory activator (Desfougères et al., 2016). Vtc4 catalyzes polyP synthesis by transferring the γ-phosphate of ATP to the growing polyP chain and releasing ADP (Hothorn et al., 2009). The transmembrane domains of the five core subunits assemble into a channel across the vacuolar membrane. In the activated complex, the exit of the Vtc4 catalytic tunnel is positioned directly above the entrance of this channel, enabling polyP synthesis on the cytosolic side of the membrane to proceed concomitantly with translocation of the polyP chain into the vacuolar lumen. VTC activity is controlled by direct binding of IPs and IPPs to its cytosolic SPX domains. The physiological IPPs 5-IP₇ and 1,5-IP₈ both activate VTC *in vitro*, with 1,5-IP₈ exhibiting a stronger effect *in vitro* (Gerasimaite et al., 2017). Consequently, the accumulation of IPPs under P_i_-replete conditions provides a direct mechanism for coupling P_i_ availability to ATP-dependent polyP synthesis and vacuolar sequestration, whereas their decline during P_i_ scarcity is expected to reduce VTC activation and preserve cytosolic P_i_ for essential metabolism (Guan et al., 2023; Pipercevic et al., 2023).

Together, these regulatory branches position 1,5-IP₈ as a coordinating signal that promotes vacuolar P_i_ storage under P_i_ sufficiency while limiting storage and activating P_i_ acquisition when intracellular P_i_ becomes scarce.

### 1.4 Process limitations and objectives of this study

Despite the growing mechanistic understanding of P_i_ sensing and polyP metabolism in *S. cerevisiae*, translating this knowledge into an efficient production process remains challenging. The P_i_ overplus response can generate a high intracellular polyP content, but its implementation requires separate biomass production, P_i_ starvation, and P_i_ refeeding stages (Christ et al., 2020).

In this study, we exploited the genetic tractability of *S. cerevisiae* to overcome the limitation of natural P_i_ overplus. Genetic interventions affecting P_i_ homeostasis and polyP metabolism were evaluated to identify a strain that combines high polyP content with accumulation under P_i_-replete conditions. The principal objective was to decouple strong polyP production from the requirement for a preceding P_i_ starvation phase, thereby simplifying the process to a single-stage fed-batch cultivation and, allowing efficient resource usage. Ultimately, by applying process-guided strain engineering, we sought to establish a production strain compatible with conventional industrial fermentation procedures, in which biomass formation and polyP accumulation occur simultaneously.

## 2. Results

To identify genetic interventions that decouple high polyP accumulation from P_i_ starvation, we followed a systematic, knowledge-driven approach by constructing a panel of 57 *S. cerevisiae* strains in the prototrophic CEN.PK113-7D background (Entian and Kötter, 2007). Because previously reported polyP phenotypes have often been characterized in auxotrophic deletion collections or heterogeneous strain backgrounds, direct comparison of their effects and assessment of their suitability for process development remained difficult (Chen et al., 2025; Hürlimann et al., 2007; Ogawa et al., 2000; Umeda et al., 2023). We therefore introduced gene deletions and overexpression cassettes *de novo* using CRISPR/Cas9, enabling their evaluation in a common haploid genetic background. Our panel covered the core PHO transcriptional circuit, P_i_ transport, IP/IPP metabolism, VTC-mediated polyP synthesis, and polyP degradation. Each strain was examined both in single-stage cultivation in P_i_-replete SD medium and under a P_i_ overplus protocol (Christ and Blank, 2019). This parallel design distinguishes interventions that primarily affected the overplus response from those that promoted polyP accumulation without a preceding starvation phase. Across both cultivation regimes, the strongest increases in polyP accumulation were associated with constitutive activation of the core PHO regulon.

### 2.1 The PHO regulon is the primary determinant of polyP accumulation

Under P_i_ overplus conditions, deletion of the negative PHO regulon component *PHO80* produced the strongest polyP accumulation among the regulatory interventions tested, while deletion of *PHO85* and overexpression of *PHO4* likewise resulted in a pronounced increase relative to the CEN.PK113-7D wild type (WT) (Fig. 2A). Conversely, deletion of the transcription factors *PHO4* or *PHO2*, or of the cyclin-dependent kinase inhibitor *PHO81*, strongly reduced polyP accumulation to a similar extent. These reciprocal phenotypes identify the Pho80–Pho85–Pho4 regulatory axis as a major determinant of the capacity to accumulate polyP following P_i_ starvation and refeeding.

**Fig. 2:**
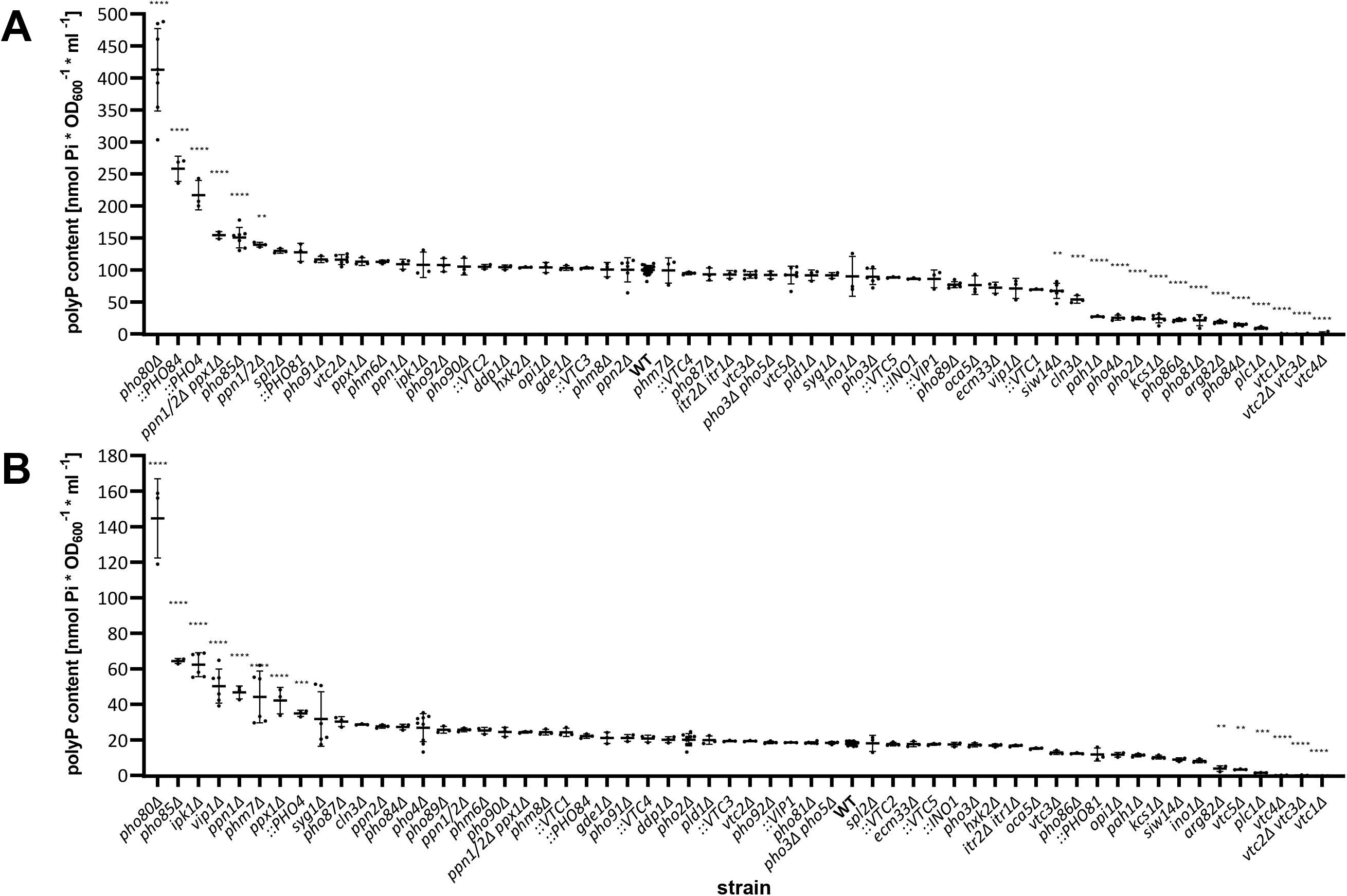
Genetic determinants of polyP accumulation during phosphate (Pi) overplus and Pi-replete cultivation. Intracellular polyP content was compared between the S. cerevisiae CEN.PK113-7D wild type (WT) 320 and derived strains carrying targeted gene deletions (Δ) or overexpression cassettes (::) following a three-stage Pi overplus protocol comprising biomass production, Pi starvation and Pi refeeding (**A**) and a single-321 stage cultivation in Pi-replete SD medium (**B**). PolyP was quantified as Pi and normalized to culture volume and biomass, expressed as nmol Pi·OD₆₀₀⁻¹·mL⁻¹. Each point represents an independent biological replicate 322 (n = 3–9 for engineered strains, n = 18 for WT). The mean is presented as black bar ± standard deviation (SD). Measurements were corrected for inter-batch variation using the WT control, and strains are ordered 323 by decreasing mean polyP content. Statistical significance was assessed by ordinary one-way ANOVA followed by Dunnett’s multiple-comparisons test against the WT. Statistical significance is indicated as follows: 324 **p < 0.01, ***p < 0.001, and ****p < 0.0001.

A similar, though not identical, pattern emerged during single-stage cultivation in P_i_-containing SD medium (Fig. 2B). *pho80Δ* again displayed the highest polyP content, and both *pho85Δ* and *PHO4* overexpression increased accumulation relative to WT. In contrast to the overplus condition, deletion of *PHO4*, *PHO2,* or *PHO81* resulted in little to no reduction in basal polyP content. Thus, constitutive PHO regulon activation strongly enhances polyP accumulation under both cultivation regimes, whereas basal synthesis during P_i_-replete growth in the absence of a PHO transcriptional response remains mostly unchanged. Nevertheless, *pho80Δ* accumulated approximately 186% more polyP following the overplus protocol than during P_i_-replete cultivation (Table 1), indicating that genetic activation alone did not account for the difference between the two processes.

**Table 1:** Comparison of polyP accumulation phenotypes that differed significantly from the wild type under phosphate (P_i_) overplus or P_i_-replete cultivation. Intracellular polyP content was compared between the S. cerevisiae CEN.PK113-7D wild type (WT) and derived strains carrying targeted gene deletions (Δ) or overexpression cassettes (::) under the P_i_ overplus and P_i_-replete cultivation protocols described for Fig. 2A and 2B. PolyP was quantified as P_i_ and normalized to culture volume and biomass, expressed as nmol P_i_·OD₆₀₀⁻¹·mL⁻¹. Data are presented as the mean ± standard deviation (SD). Values not distinguishable from the blank are reported as not detected (n.d.). Standard deviations were rounded to two significant figures, and means were rounded to the corresponding decimal place. Statistical significance (St. sig.) was assessed by ordinary one-way ANOVA followed by Dunnett’s multiple-comparisons test against the WT. Statistical significance is indicated as follows: **p < 0.01, ***p < 0.001, and ****p < 0.0001.

| **Strain name** | **PolyP content under P<sub>i</sub> overplus conditions** [nmol P<sub>i</sub>·OD<sub>600</sub><sup>−1</sup>·mL<sup>−1</sup>] | **SD** | **St. sig.** | **PolyP content under P<sub>i</sub>-replete conditions** [nmol P<sub>i</sub>·OD<sub>600</sub><sup>−1</sup>·mL<sup>−1</sup>] | **SD** | **St. sig.** |
|---|---|---|---|---|---|---|
| *pho80Δ* | 413 | 64 | **** | 145 | 22 | **** |
| *::PHO84* | 258 | 20 | **** | 22.1 | 1.2 |  |
| *::PHO4* | 217 | 23 | **** | 34.9 | 1.8 | *** |
| *ppn1Δ ppn2Δ ppx1Δ* | 154.4 | 5.5 | **** | 24.3 | 0.5 |  |
| *pho85Δ* | 150 | 16 | **** | 64.4 | 1.4 | **** |
| *ppn1Δ ppn2Δ* | 139.4 | 3.7 | ** | 25.6 | 1.1 |  |
| *ppx1Δ* | 113.1 | 6.0 |  | 42.1 | 7.5 | **** |
| *ppn1Δ* | 108.8 | 7.8 |  | 46.7 | 3.6 | **** |
| *ipk1Δ* | 108 | 20 |  | 62.3 | 6.7 | **** |
| **WT** | 100.0 | 7.0 |  | 18.0 | 1.0 |  |
| *phm7Δ* | 99 | 20 |  | 44 | 15 | **** |
| *vtc5Δ* | 92 | 14 |  | 3.3 | 0.2 | ** |
| *vip1Δ* | 71 | 16 |  | 50.2 | 9.5 | **** |
| *siw14Δ* | 67 | 12 | ** | 8.8 | 1.0 |  |
| *cln3Δ* | 54.0 | 6.2 | *** | 28.6 | 0.4 |  |
| *pah1Δ* | 27.1 | 1.0 | **** | 11.3 | 0.9 |  |
| *pho4Δ* | 25.3 | 3.9 | **** | 26.8 | 7.9 |  |
| *pho2Δ* | 24.2 | 2.0 | **** | 20.0 | 3.5 |  |
| *kcs1Δ* | 23.8 | 6.5 | **** | 10.2 | 1.1 |  |
| *pho86Δ* | 21.9 | 2.5 | **** | 12.3 | 0.4 |  |
| *pho81Δ* | 21.3 | 8.6 | **** | 18.4 | 0.6 |  |
| *arg82Δ* | 18.7 | 2.6 | **** | 3.8 | 1.6 | ** |
| *pho84Δ* | 14.3 | 1.9 | **** | 27.2 | 1.6 |  |
| *plc1Δ* | 9.4 | 2.0 | **** | 1.5 | 0.1 | *** |
| *vtc1Δ* | n.d. |  | **** | n.d. |  | **** |
| *vtc2Δ vtc3Δ* | n.d. |  | **** | n.d. |  | **** |
| *vtc4Δ* | n.d. |  | **** | n.d. |  | **** |

These phenotypes are consistent with several previous reports linking constitutive PHO regulon activation to increased polyP accumulation in other *S. cerevisiae* strain backgrounds (Hürlimann et al., 2007; Umeda et al., 2023).

### 2.2 Polyphosphatase deletions moderately increase polyP accumulation

Interventions targeting polyP degradation produced smaller and more condition-dependent effects compared to constitutive PHO regulon activation. Under P_i_ overplus conditions, combined deletion of *PPN1*, *PPN2*, and *PPX1* increased polyP accumulation by approximately 50% relative to WT (Fig. 2A). During single-stage cultivation, the same strain exhibited an increase of approximately 35%, which did not reach statistical significance (Fig. 2B). Conversely, the individual *ppn1Δ* and *ppx1Δ* strains accumulated approximately twice as much polyP as the WT during single-stage cultivation.

Increased polyP accumulation following deletion of *PPN1* or *PPX1* is consistent with previous observations in another *S. cerevisiae* background (Umeda et al., 2023). However, the effects of the individual deletions were not additive in the triple mutant under P_i_-replete growth, indicating that the contribution of the different polyphosphatases cannot be inferred simply from their combination. Since these phenotypes were identified within the broad strain screen and displayed considerable differences between the two tested conditions, the relative effects of the individual and combined deletions under Pi-replete conditions require direct side-by-side validation. Nevertheless, none of the polyphosphatase mutants approached the polyP accumulation achieved by *pho80Δ*, suggesting that limiting polyP degradation alone was insufficient to generate the desired production phenotype.

### 2.3 IP metabolism affects polyP accumulation in a condition-dependent manner

Disruption of early steps in IP and IPP biosynthesis strongly impaired polyP accumulation. Deletion of *PLC1*, *ARG82,* or *KCS1* reduced polyP under both cultivation regimes, reproducing the phenotypes previously reported by Auesukaree et al. (2005). Because these mutations are associated with constitutive PHO regulon activation, their low polyP content further indicates that transcriptional activation of the PHO response is not sufficient for polyP accumulation when downstream IPP synthesis is compromised. In contrast, deletion of *IPK1* or *VIP1* did not reduce polyP accumulation during the overplus protocol and significantly increased it during single-stage cultivation, with both mutants ranking among the strongest non-PHO regulon candidates under this condition. Elevated polyP accumulation in an *ipk1Δ* strain was also reported by Lonetti et al. (2011). Similarly, increased polyP accumulation following deletion of *VIP1* was recently observed in the BY4741 background (Chen et al., 2025).

Deletion of the IPP phosphatases *SIW14*, *DDP1,* or *OCA5* produced smaller or condition-dependent changes and did not generate a similarly robust production phenotype. Collectively, these results distinguish the requirement for the upstream formation of highly phosphorylated inositol species from the more variable consequences of perturbing their downstream interconversion.

### 2.4 PolyP accumulation depends on the VTC complex

The deletion of VTC complex components resulted in some of the strongest loss-of-function phenotypes in the screening. Deletion of *VTC1,* coding for a structural part of the polyP-synthesizing VTC complex, or *VTC4,* coding for the catalytic polymerase subunit of the VTC complex, resulted in no detectable polyP under either cultivation condition, consistent with their established central roles in vacuolar polyP synthesis (Ogawa et al., 2000; Hothorn et al., 2009). Individual deletion of *VTC2* or *VTC3* caused small changes that did not reach the significance threshold, whereas simultaneous deletion of both genes had no detectable polyP in either protocol. This pattern supports the functional overlap of Vtc2 and Vtc3 as alternative components of related VTC assemblies (Pipercevic et al., 2023). In contrast, deletion of *VTC5* markedly impaired polyP accumulation during single-stage P_i_-replete cultivation but did not produce a comparable defect following P_i_ overplus. Thus, although Vtc5 is an established positive regulator of VTC activity under standard growth conditions (Desfougères et al., 2016), it appears dispensable for the strong activation of polyP synthesis during the P_i_ overplus response. Moreover, overexpression of individual VTC subunits did not increase polyP accumulation (Umeda et al., 2023). These findings confirm the VTC complex as the principal polyP synthesis machinery under both cultivation regimes while revealing a condition-specific requirement for its regulatory subunit Vtc5.

### 2.5 Pho84-dependent P_i_ uptake is required for the P_i_ overplus response

Because VTC-mediated polyP synthesis ultimately depends on the supply of cytosolic P_i_, we next examined the contribution of P_i_ uptake. Deletion of the high-affinity P_i_ transporter *PHO84* strongly reduced polyP accumulation following P_i_ overplus, recapitulating the low-polyP and low-uptake phenotypes previously reported for *pho84Δ* cells under P_i_ overplus conditions (Ogawa et al., 2000). Deletion of *PHO86*, which is required for efficient trafficking of Pho84 from the endoplasmic reticulum to the plasma membrane, produced a similarly pronounced reduction (Lau et al., 2000). By contrast, neither deletion caused a comparable loss of polyP during P_i_-replete cultivation, and none of the individual P_i_ transporter deletions (*PHO87*, *PHO89, PHO90*) significantly affected polyP accumulation under this condition. The absence of a P_i_-replete phenotype in *pho84Δ* agrees with previous observations that intracellular P_i_ and polyP remain approximately unchanged despite constitutive *PHO5* activation (Auesukaree et al., 2005), as well as with a recent analysis of P_i_ transporter mutants under high-P_i_ conditions (Ledova et al., 2025), and suggests substantial functional redundancy, with the remaining uptake systems compensating for individual transporter deletions when extracellular P_i_ is abundant. This condition-dependent phenotype indicates that Pho84-mediated uptake becomes particularly important under P_i_ overplus, whereas alternative P_i_ transporters can sustain polyP synthesis under P_i_-replete conditions. Together, these results identify the Pho84–Pho86 transport system as the principal route supplying P_i_ for the P_i_ overplus response.

### 2.6 *PHM7* deletion produces a condition-dependent polyP phenotype

Finally, deletion of *PHM7* produced a condition-specific phenotype that distinguished it from the other strains examined. Whereas *phm7Δ* did not significantly affect polyP accumulation following P_i_ overplus, it increased polyP content during P_i_-replete cultivation. Phm7 remains a poorly characterized PHO-associated protein, although it has been implicated in supporting P_i_ uptake and P_i_ overplus accumulation. Notably, this phenotype contrasts with the reduced P_i_ uptake and polyP accumulation previously reported for *phm7Δ* cells during YPD cultivation (Kulakovskaya et al., 2023). Together with the substantial biological variability observed in this measurement, this discrepancy suggests that the influence of *PHM7* on polyP accumulation may depend strongly on the strain background or cultivation conditions. The increased P_i_-replete polyP phenotype should therefore be regarded as a preliminary observation requiring independent validation.

### 2.7 *PHO80* deletion enables high polyP accumulation without P_i_ starvation

The initial screening experiments revealed a 186% difference in the polyP content of *S. cerevisiae* CEN.PK113-7D *pho80Δ* (hereafter referred to as *pho80Δ*) cells between the P_i_ overplus and P_i_-replete protocols. To determine whether this difference resulted specifically from P_i_ starvation, we compared CEN.PK113-7D *pho80Δ* with the parental WT and the high-polyP reference strain *S. cerevisiae* VH2.200, identified by Christ and Blank (2019), using the same protocols either including or omitting the starvation phase (Fig. 3A). Following starvation and refeeding, VH2.200 and *pho80Δ* accumulated 23.0% and 21.6% polyP as KPO_3_ in cell dry weight (CDW), respectively, whereas the parental CEN.PK113-7D strain accumulated 1.9%. Omitting P_i_ starvation strongly reduced polyP accumulation in both reference strains. In contrast, the polyP content of *pho80Δ* remained at 20.6%, demonstrating that deletion of *PHO80* eliminates the requirement for starvation-dependent induction of high polyP accumulation. The lower polyP content of *pho80Δ* observed during the initial single-stage screening experiment therefore cannot be attributed to the absence of P_i_ starvation, indicating that additional differences between the cultivation protocols influence the attainable polyP content.

**Fig. 3.**
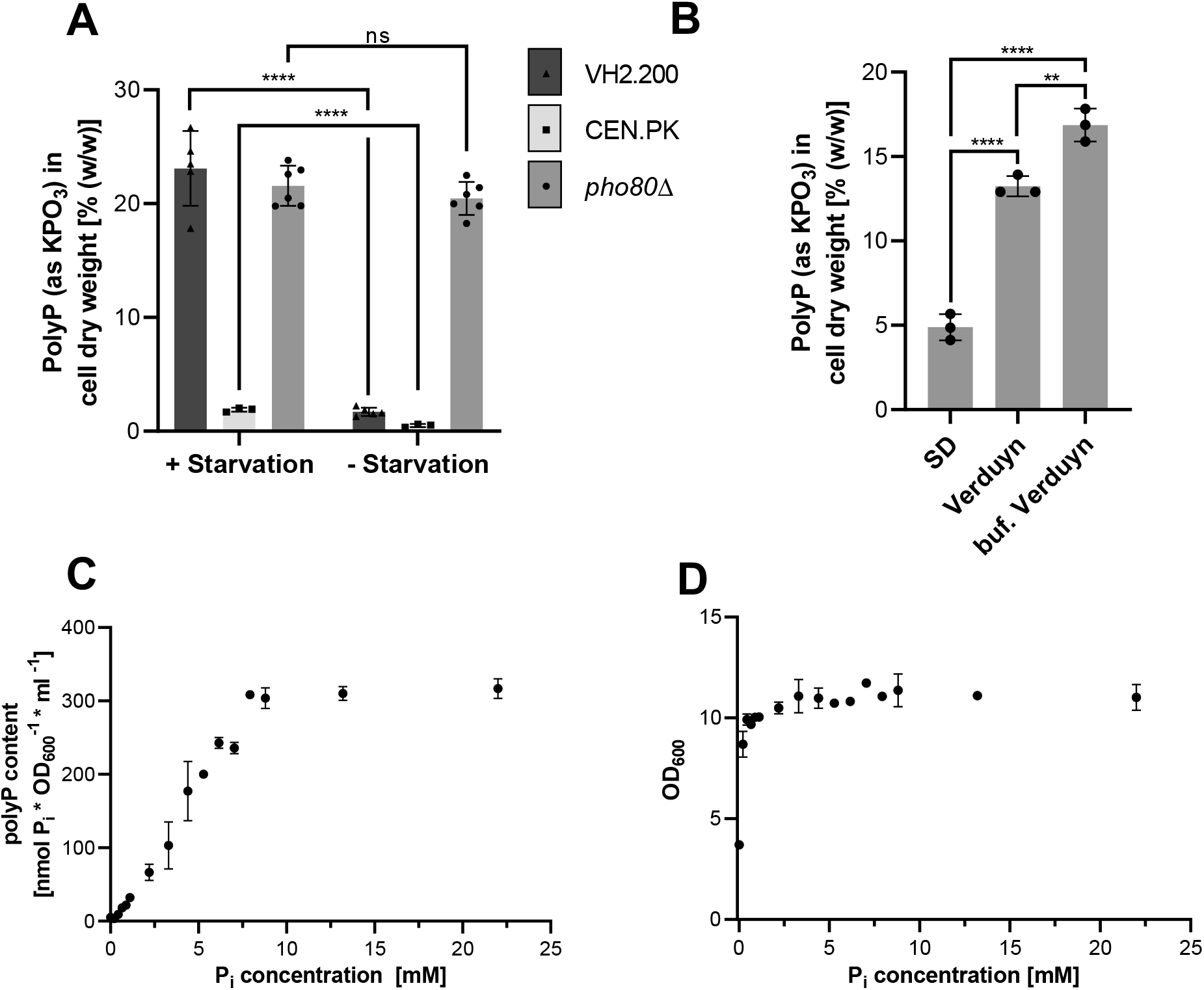
PHO80 deletion enables starvation-independent polyP accumulation, while medium composition, pH, and P_i_ availability determine the attainable polyP content. (A) PolyP content of S. cerevisiae VH2.200, CEN.PK113-7D wild type and CEN.PK113-7D pho80Δ following an otherwise equivalent P_i_ overplus protocol either including (+ starvation) or omitting (− starvation) the 6 h cultivation phase in P_i_-free medium. PolyP is expressed as KPO₃ equivalents relative to cell dry weight. (B) PolyP content of CEN.PK113-7D pho80Δ after cultivation in SD medium, unbuffered Verduyn medium or Verduyn medium buffered to pH 5.2 with potassium hydrogen phthalate. (C) PolyP content of CEN.PK113-7D pho80Δ cultivated in buffered Verduyn medium containing different initial P_i_ concentrations. PolyP was quantified as released P_i_ and is expressed as nmol P_i_·OD₆₀₀⁻¹·mL⁻¹. (D) Final OD₆₀₀ values of the cultures shown in panel C. In panels A and B, individual symbols represent independent biological replicates and columns show mean ± SD (n = 3–6 in A and n = 3 in B). Data in panels C and D are presented as mean ± SD from three biological replicates (n = 3). Statistical significance in panel A was assessed by two-way ANOVA followed by Tukey’s multiple-comparisons test; only the within-strain comparisons between cultivation with and without starvation are shown. Statistical significance in panel B was assessed by one-way ANOVA followed by Tukey’s multiple-comparisons test. Statistical significance is indicated as follows: ns, not significant; **p < 0.01 and ****p < 0.0001.

### 2.8 Medium composition and pH stabilization increase polyP accumulation in *pho80Δ*

Having established that P_i_ starvation itself was not required for high polyP accumulation in *pho80Δ*, we next investigated whether the lower polyP content observed during single-stage SD cultivation resulted from the medium composition. Previous observations indicated that P_i_ can become limiting in SD medium and that the culture pH decreased from 5.4 to approximately 2.8 during cultivation. In SD medium, *pho80Δ* accumulated 4.9% polyP (Fig. 3B). Cultivation in Verduyn medium, which contained a higher initial P_i_ concentration, significantly increased the polyP content to 13.2%. Stabilizing the Verduyn medium at pH 5.2 with potassium hydrogen phthalate produced a further significant increase to 16.8%. These results show that the comparatively low polyP content observed in the initial P_i_-replete screen is attributable to media effects and culture acidification rather than to the absence of a starvation phase.

### 2.9 PolyP accumulation requires higher P_i_ availability than biomass formation

To determine how extracellular P_i_ availability constrains polyP production, CEN.PK113-7D *pho80Δ* was cultivated in buffered Verduyn medium containing different initial P_i_ concentrations (Fig. 3C and D). PolyP content in the biomass remained at a plateau between 7.9 mM and 22 mM initial P_i_, indicating that additional P_i_ no longer increased storage above this range. Below 7.9 mM, polyP content decreased approximately linearly with decreasing initial P_i_ concentration. At initial concentrations below 8.8 mM, residual P_i_ in the culture supernatant was below the quantification limit after cultivation, indicating near-complete consumption of the supplied P_i_ (Supplementary Fig. S2). In contrast to polyP accumulation, biomass formation remained largely unchanged until the initial P_i_ concentration fell below 0.44 mM. Thus, P_i_ concentrations that were sufficient to support near-maximal growth were insufficient to sustain high polyP accumulation. These findings demonstrate that polyP formation represents a storage response that occurs only after the cellular P_i_ requirements for biomass formation have been met.

### 2.10 *PHO80* deletion constitutively induces the *PHO5* promoter

To evaluate how the *PHO80* deletion influenced the activation of the PHO regulon in comparison to WT, an expression cassette containing superfolder green fluorescent protein (sfGFP) under the control of the *PHO5* promoter was introduced into the WT and the CEN.PK113-7D *pho80Δ* strain, as previously employed to monitor P_i_ starvation (Deitert et al., 2025; Lam et al., 2008). Both strains were cultivated in medium containing different concentrations of P_i_, and biomass formation and GFP fluorescence were monitored online during the cultivation (Fig. 4A and B). In the WT, GFP expression is triggered as a consequence of P_i_ consumption during growth and is observed when cultivated at a concentration below 3.3 mM. Induction of GFP expression occurred over time as the P_i_ concentration decreased below the induction threshold for *PHO5* expression and occurred at the lowest concentrations first. Expression strength also correlated with lower P_i_ concentration, except when 0 mM P_i_ was supplied, probably due to impaired growth and cellular maintenance.

**Fig. 4:**
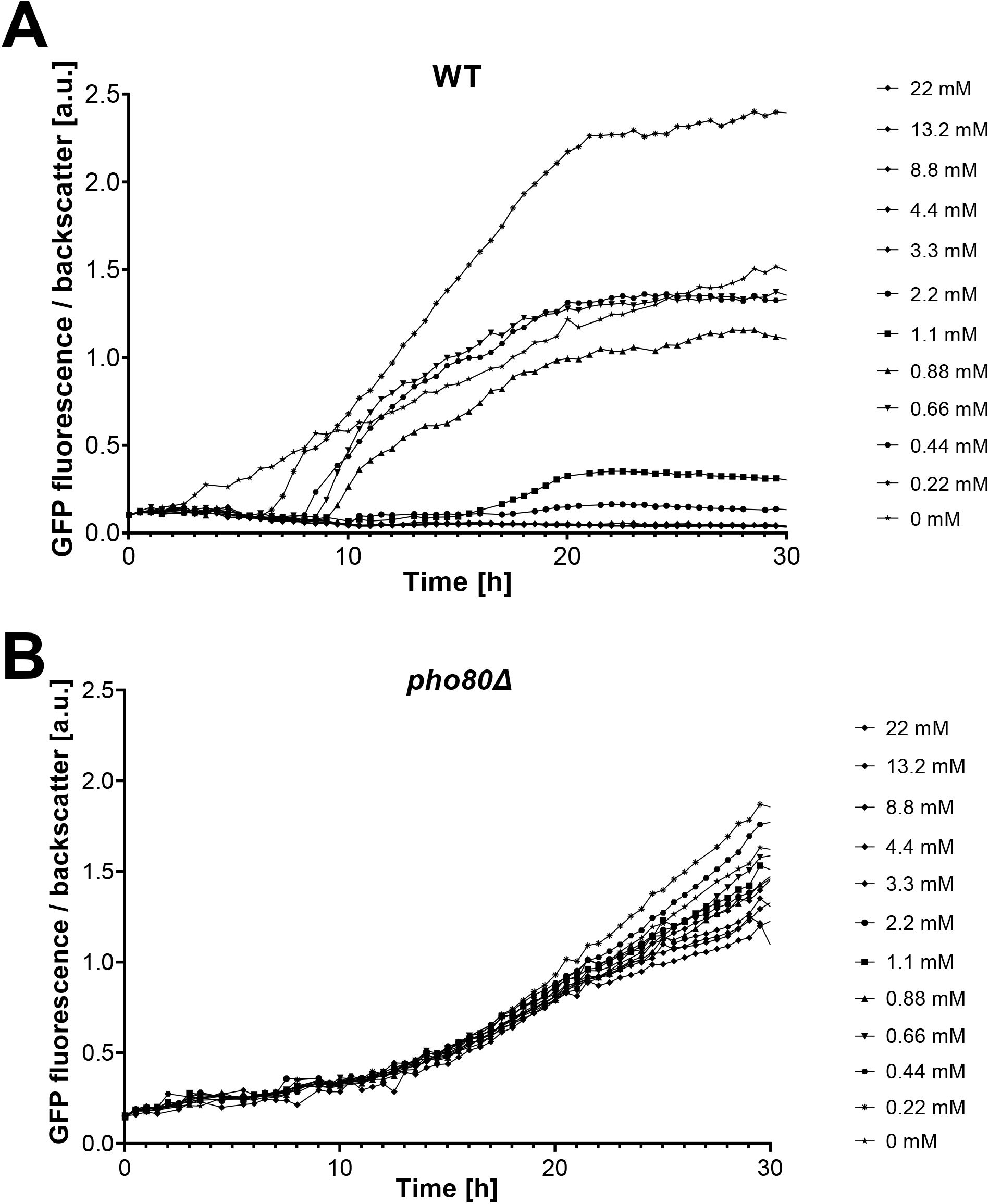
PHO80 deletion induces PHO regulon activity independent of the initial P_i_ concentration. Time courses of GFP fluorescence normalized to the biomass signal in CEN.PK113-7D strains carrying an integrated sfGFP reporter under the control of the PHO5 promoter. Reporter activity was monitored in the wild type (**WT**) and **pho80Δ** during cultivation in buffered Verduyn medium containing the indicated initial P_i_ concentrations ranging from 0 to 22 mM. Fluorescence is presented in arbitrary units. Data represent the mean of two independent biological replicates; error bars were omitted for visual clarity, and single replicates are shown in Supplementary Fig. S3.

In the *pho80Δ* strain, induction of GFP expression is seen independent of P_i_ concentration from the start of the cultivation. Expression strength is slightly elevated for the lower concentration conditions at the end of cultivation, possibly reflecting further induction of the PHO regulon by declining IPP levels (Auesukaree et al., 2005) or accumulation of AICAR (Kim et al., 2023).

### 2.11 Electron microscopy reveals enlarged vacuoles and vacuolar phosphorus enrichment in *pho80Δ*

Initial observations of *pho80Δ* cells and of WT cells after P_i_ starvation and refeeding suggested increased vacuolar size. To examine vacuolar morphology and phosphorus localization, we compared CEN.PK113-7D WT, the polyP-deficient *vtc4Δ* strain, and the polyP-accumulating *pho80Δ* strain by transmission electron microscopy (TEM) and scanning electron microscopy (SEM) coupled with energy-dispersive X-ray spectroscopy (EDX) (Fig. 5). In the examined TEM sections, larger vacuoles were observed in *pho80Δ* compared to WT and *vtc4Δ* (Fig. 5A). In the SEM images, *pho80Δ* vacuoles appeared bright, whereas those in WT and *vtc4Δ* were grayish and dark, respectively (Fig. 5B). This contrast may reflect differences in vacuolar composition and heavy-metal staining. The phosphorus K_α_-signal maps showed localized enrichment at vacuolar positions in *pho80Δ*, as indicated by the overlays with the SEM images (Fig. 5C and D). Localized vacuolar phosphorus signals were also visible in WT, whereas no clear vacuolar enrichment was apparent in *vtc4Δ*.

**Fig. 5:**
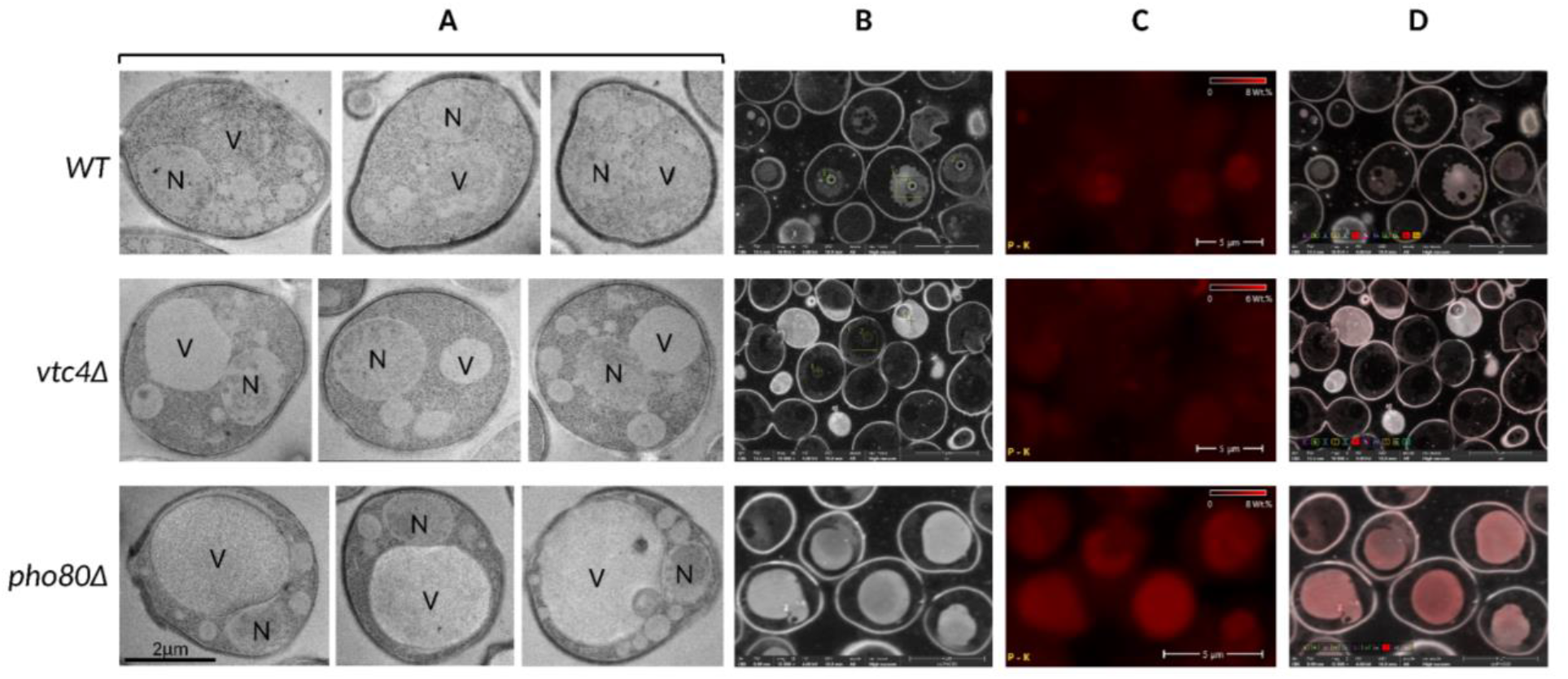
TEM, SEM, and EDX phosphorus mapping of CEN.PK113-7D wild type (WT), vtc4Δ, and pho80Δ. Cells were cultivated in buffered Verduyn medium and harvested at an OD_600_ of 1. (**A**) Three representative TEM images per strain; V, vacuole; N, nucleus. (**B**) SEM images. (**C**) Corresponding EDX maps of the phosphorus K_α_ signal, shown in red. (**D**) Overlays of the SEM images in B and the phosphorus maps in C. Rows show WT, vtc4Δ, and pho80Δ from top to bottom.

Together with the polyP measurements, the vacuolar phosphorus enrichment in *pho80Δ* and the lack of clear enrichment in the polyP-deficient *vtc4Δ* control support vacuolar storage of the accumulated polyP, which is also backed by literature (Gerasimaitė et al., 2014; Saito et al., 2005; Urech et al., 1978). The enlarged vacuolar profiles suggest that enhanced storage is accompanied by altered vacuolar morphology during growth without prior P_i_ starvation. Since EDX detects elemental phosphorus, the maps provide no polyP-specific measurement.

### 2.12 *PHO80* deletion enables high-titer polyP production under industrially relevant fed-batch conditions

The preceding experiments identified CEN.PK113-7D *pho80Δ* as a promising production strain because it accumulated polyP during exponential growth in buffered, P_i_-replete medium while consuming nearly all supplied P_i_. Its maximum specific growth rate was determined as 0.31 ±0.02 h⁻¹, corresponding to a 23% reduction relative to the parental strain (0.40 ±0.03 h⁻¹) (Supplementary Fig. S1), but remaining above the approximate 0.18 h⁻¹ considered sufficient for the oxygen transfer rates achievable in industrial-scale fermentation (Attfield, 2022; Van Hoek et al., 2000). Based on their elevated polyP accumulation, *pho80Δ* and *pho85Δ*, which disrupt the Pho80–Pho85 cyclin–cyclin-dependent kinase complex, were evaluated in aerobic 0.2 L exponential fed-batch cultivations (Fig. 6 and Table 2).

**Fig. 6:**
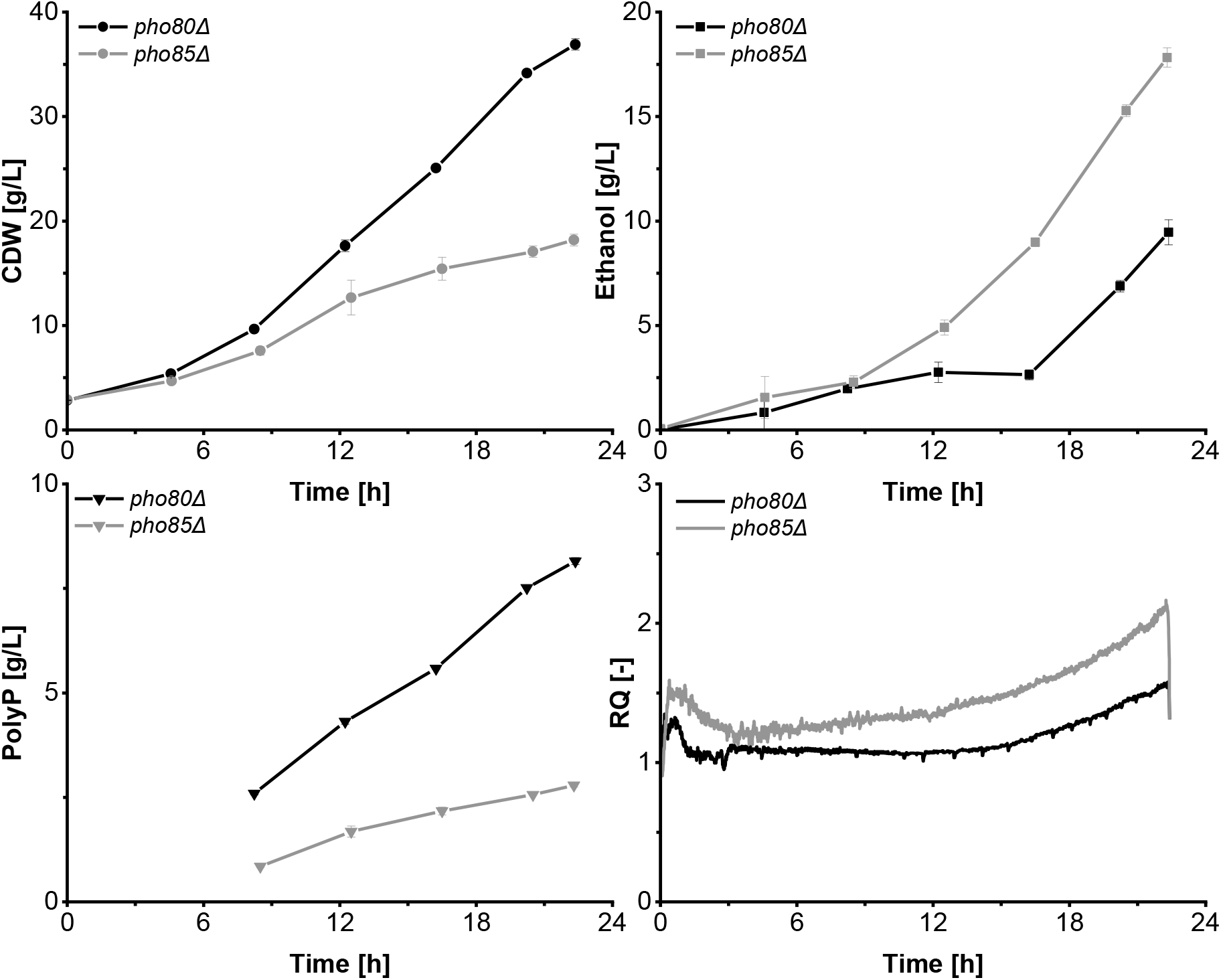
Aerobic fed-batch production of polyP by CEN.PK113-7D pho80Δ and pho85Δ. Time courses of cell dry weight (CDW), ethanol concentration, polyP titer, and respiratory quotient (RQ) during 0.2 L exponential fed-batch cultivation. polyP titers are expressed as KPO₃ equivalents. The RQ was calculated as the molar ratio of the carbon dioxide evolution rate to the oxygen uptake rate. Lines connect consecutive measurements and serve as visual guides. Data are presented as the mean ± SD of two independent cultivations (n = 2).

**Table 2:** Key performance indicators of aerobic fed-batch cultivation of CEN.PK113-7D pho80Δ and pho85Δ. Data are presented as the mean ± SD from two independent cultivations (n=2).

| **Performance indicator** | ***pho80Δ*** | ***pho85Δ*** | **Unit** |
|---|---|---|---|
| Fermentation time | 22.4 | 22.3 | h |
| Cell dry weight at harvest | 36.9 ±0.6 | 18.2 ±0.6 | g/L |
| PolyP content at harvest (KPO<sub>3</sub> in CDW) | 22.1 ±0.1 | 15.3 ±0.3 | % (w/w) |
| PolyP content at harvest (P in CDW) | 5.8 ±0.0 | 4.0 ±0.1 | % (w/w) |
| PolyP titer at harvest (as KPO<sub>3</sub>) | 8.2 ±0.1 | 2.8 ±0.0 | g/L |
| PolyP titer at harvest (as P) | 2.1 ±0.0 | 0.7 ±0.0 | g/L |
| Biomass yield on glucose (CDW/glucose) | 0.46 ±0.01 | 0.22 ±0.01 | g/g |
| Final ethanol concentration | 9.46 ±0.60 | 17.82 ±0.50 | g/L |
| Average specific growth rate (µ) | 0.146 ±0.002 | 0.113 ±0.007 | 1/h |
| Biomass space time yield (CDW/V/t) | 1.52 ±0.03 | 0.69 ±0.04 | g/L/h |
| PolyP space time yield (w/V/t) | 0.36 ±0.00 | 0.13 ±0.00 | g/L/h |

The *pho80Δ* strain initially followed the exponential feed profile. After approximately 12 h, its respiratory quotient increased, and more ethanol began to accumulate, indicating the onset of respiro-fermentative overflow metabolism. Nevertheless, biomass and polyP continued to increase until harvest. By contrast, *pho85Δ* grew more slowly than expected based on the feed profile and showed higher ethanol accumulation, consistent with the glucose supply exceeding its respiratory growth capacity and being partly diverted to ethanol formation. No residual glucose was detected at any sampling point in either cultivation.

At harvest, *pho80Δ* reached a polyP titer of 8.2 g/L, approximately threefold higher than *pho85Δ*. This advantage resulted from the combination of greater biomass formation and a higher intracellular polyP content, while ethanol formation remained lower. The complete set of fermentation performance indicators is summarized in Table 2. Together, these findings establish *PHO80* deletion as a suitable basis for producing polyP under a conventional aerobic fed-batch regime without a preceding P_i_ starvation phase.

## 3. Discussion

The principal contribution of this study is the conversion of a low-polyP accumulating laboratory strain into a production strain capable of accumulating high polyP levels independent of prior P_i_ starvation. A single genetic intervention, deletion of *PHO80*, enabled CEN.PK113-7D to retain approximately 20% polyP in cell dry weight when the starvation phase was omitted, whereas the parental strain and the reference strain VH2.200 remained dependent on starvation and refeeding for strong accumulation. Importantly, this phenotype was transferred from microscale cultivation to aerobic fed-batch production, in which *pho80Δ* reached a polyP content of 22.1% of CDW, a titer of 8.2 g/L and a volumetric productivity of 0.36 g/L/h, expressed as KPO_3_ equivalents. While *PHO80* has previously been associated with P_i_ homeostasis and polyP accumulation, the significance of our study lies in using its deletion to eliminate the overplus requirement, permitting polyP accumulation during biomass formation. This trait establishes compatibility with an exponential fed-batch operation, which is a prerequisite for industrial production with *S. cerevisiae*. The *PHO80* deletion therefore converts a transient physiological P_i_ overplus response into a constitutive production phenotype.

The broader strain screening identifies the Pho80–Pho85–Pho4 regulatory axis as the most effective engineering layer for increasing polyP accumulation. Deletion of *PHO80* or *PHO85* and overexpression of *PHO4* resulted in substantially stronger effects than overexpression of individual VTC components or deletion of polyphosphatases. This indicates that polyP production is primarily defined by coordinated pathway regulation rather than by the abundance of an individual VTC subunit or by degradation rate. Manipulation of an upstream regulatory component can simultaneously increase the capacity for P_i_ acquisition, scavenging, and storage, whereas increasing one VTC component might be ineffective because polymerase activity depends on correct complex assembly, subunit stoichiometry and post-translational activation by IPPs. The comparison between *pho80Δ* and *pho85Δ* further illustrates the importance of target specificity. Both deletions activate the PHO regulon, but *pho85Δ* exhibited slower growth, lower biomass yield, and greater ethanol formation during fed-batch cultivation. This stronger fitness impairment may reflect the broader pleiotropic function of Pho85. Whereas deletion of *PHO80* specifically disrupts the Pho80–Pho85 complex, deleting *PHO85* eliminates Pho85-dependent kinase activity associated with multiple cyclin partners involved in nutrient signaling, metabolism, and cell-cycle regulation (Carroll and O’Shea, 2002). Targeting the PHO-specific cyclin Pho80 consequently provides a more favorable balance between pathway activation and host fitness.

Together, the observed phenotypes support a two-input model for efficient polyP accumulation. First, derepression of the PHO regulon provides the cellular capacity for P_i_ acquisition and storage. Second, a functional IPP–VTC branch is required to translate this capacity into polyP synthesis. This interpretation is supported by the low polyP contents of *plc1Δ*, *arg82Δ* and *kcs1Δ*, despite the constitutive PHO regulon activation expected in these backgrounds (Auesukaree et al., 2005), and by the absence of detectable polyP following deletion of *VTC1*, *VTC4,* or both *VTC2* and *VTC3*. Conversely, constitutive activity of the *PHO5* promoter in *pho80Δ* confirms that *PHO80* deletion bypasses P_i_-dependent repression of a central PHO-regulated promoter. These findings indicate that transcriptional activation is necessary for maximal polyP accumulation but is not sufficient when synthesis of the IPP signals required for VTC activation or the VTC polymerase itself is disrupted.

This hypothesis also provides a mechanistic explanation for the starvation-independent phenotype of *pho80Δ*. In the WT, activation of the PHO regulon normally accompanies declining P_i_ and 1,5-IP₈ concentrations, whereas strong VTC activation is favored under P_i_-replete conditions in which activating IPPs are available (Chabert et al., 2023; Pipercevic et al., 2023). P_i_ overplus may transiently bridge these states because proteins induced during starvation remain available after P_i_ refeeding restores the substrate and signaling conditions required for VTC activity. Deletion of *PHO80* bypasses the repression of the transcriptional branch, while cultivation under P_i_-replete conditions may preserve ATP, P_i_, and IPP conditions permissive for VTC activation. Functions that are normally separated temporally could therefore operate simultaneously, converting transient overplus accumulation into sustained polyP production during growth. This interpretation is consistent with the genetic and reporter data but remains a working hypothesis as IPP concentrations were not quantified.

The increased P_i_-replete polyP accumulation observed in *ipk1Δ* and *vip1Δ* further illustrates the complexity of the IPP network. Reported polyP phenotypes for these deletions differ among studies, including normal and elevated accumulation depending on the strain background and cultivation conditions (Auesukaree et al., 2005; Chen et al., 2025; Lonetti et al., 2011). Because deletion of *IPK1* or *VIP1* alters several IPP species, these phenotypes cannot be attributed unambiguously to the accumulation or depletion of a single isomer. The smaller and condition-dependent effects of deleting *SIW14*, *DDP1*, or *OCA5* reinforce this limitation of genetic inference.

The loss of detectable polyP in *vtc2Δ vtc3Δ*, despite the weaker phenotypes of the individual deletions, supports the function of Vtc2 and Vtc3 as overlapping alternative components of VTC assemblies. By contrast, Vtc5 was required during P_i_-replete cultivation but dispensable for strong accumulation following P_i_ overplus, suggesting that the regulatory requirements for VTC activation differ between steady P_i_ supply and acute refeeding. P_i_ transport showed a similar distinction. The strong effects of *pho84Δ* and *pho86Δ* during P_i_ overplus indicate that rapid Pho84-dependent uptake is important when P_i_-starved cells encounter renewed P_i_ availability. However, no individual plasma-membrane transporter deletion significantly affected polyP under P_i_-replete conditions, indicating compensation by the remaining uptake systems when extracellular P_i_ is abundant. Individual transporter engineering therefore does not appear to address the principal limitation under these conditions. Polyphosphatase deletions produced moderate and non-additive effects, suggesting compensatory turnover pathways or altered localization. Finally, the variable and literature-inconsistent phenotype of *phm7Δ* makes *PHM7* a secondary candidate that requires independent validation rather than a basis for a mechanistic conclusion (Kulakovskaya et al., 2023).

Electron microscopy provides a cellular context for the production phenotype of *pho80Δ* (Fig. 5). Vacuolar phosphorus enrichment, together with the polyP measurements and the lack of enrichment in the polyP-deficient *vtc4Δ* control, supports vacuolar storage of the accumulated polyP (Saito et al., 2005). The enlarged vacuolar profiles suggest that increased storage is accompanied by altered vacuolar morphology during growth without prior P_i_ starvation. This vacuolar enlargement could accommodate greater storage.

The production phenotype of *pho80Δ* remained strongly dependent on the cultivation environment. The comparatively low polyP content obtained in the initial screening using SD medium under P_i_-replete conditions therefore underestimated the capacity of the strain. The direct comparison of cultivation with and without starvation demonstrates starvation independence, whereas the increase from SD to Verduyn medium shows that medium composition strongly affects the attainable polyP content. The comparison between buffered and unbuffered Verduyn medium directly establishes a beneficial effect of pH stabilization.

The P_i_ titration experiment provides an initial quantitative basis for medium and feed design. Biomass formation remained largely unaffected at P_i_ concentrations that already restricted polyP accumulation, demonstrating that growth is a poor indicator of whether sufficient P_i_ has been supplied for polyP production. Under the tested conditions, an initial P_i_ concentration of approximately 7.9 mM marked the onset of a plateau in biomass-normalized polyP accumulation. This value should be regarded as a process-specific starting point rather than a universal physiological threshold because P_i_ demand will depend on biomass concentration, medium composition, and cultivation mode. At lower initial concentrations, residual P_i_ fell below the analytical quantification limit, supporting near-complete P_i_ consumption. Future process optimization must therefore balance cellular polyP content and P_i_ recovery against residual P_i_ and the total P_i_ input. The data suggest that a well-balanced process is able to use most of the supplied P_i_. This is highly relevant, particularly if industrial side streams are targeted for P_i_ recovery (Fees et al., 2022; Herrmann et al., 2023).

The fed-batch fermentations demonstrate that the *pho80Δ* phenotype is compatible with an industrially relevant operating principle. Its advantage over *pho85Δ* did not solely result from cellular polyP content but from the combination of strong accumulation with substantially greater biomass formation. The average specific growth rate of *pho80Δ* initially remained close to that imposed by the exponential feed (µ_set_ = 0.15 h^−1^), whereas the slower growth of *pho85Δ* caused substrate supply to exceed its metabolic capacity. In *pho80Δ,* the increase in respiratory quotient and ethanol accumulation after approximately 12 h similarly indicate that the predefined exponential feed eventually exceeded respiratory processing capacity. This behavior represents a clear optimization opportunity: reducing the feed rate or implementing respiratory-quotient-, dissolved-oxygen-, or ethanol-responsive feeding could limit overflow metabolism and redirect more carbon toward biomass and polyP production. It would also be interesting to determine whether the later deviation results from declining growth capacity or depletion of another nutrient. Nevertheless, the conducted fed-batch fermentations demonstrate fundamental proof of compatibility with industrially relevant production. Comparisons with existing biological processes for P removal should distinguish cellular polyP content, volumetric titer, productivity, P_i_ recovery, cultivation duration, and the number of process stages.

### 3.1 Conclusion

Overall, the *PHO80* deletion provides a targeted regulatory intervention that combines constitutive expression of P_i_-responsive functions with P_i_-replete VTC-mediated storage, converting the native P_i_ overplus response into a fermentation-compatible production phenotype. The aerobic fed-batch cultivations demonstrate that CEN.PK113-7D *pho80Δ* combines high cellular polyP accumulation with biomass formation under conventional fermentation conditions. Eliminating P_i_ starvation substantially simplifies the polyP production process by unifying cultivation stages, permitting biomass and polyP formation within one fermentation. Together with the biochemical measurements, electron microscopy supports vacuolar polyP storage in *pho80Δ* and links enhanced accumulation to altered vacuolar morphology. The unification of biomass formation and polyP accumulation represents a prime example of process-guided strain engineering, leveraging metabolic and strain engineering to not only enhance titers and yields, but to increase processability (Karmainski et al., 2026). In our case, the engineered strain approaches the cellular polyP contents previously obtained through P_i_ overplus while considerably simplifying the production process. In conclusion, our approach advances biological polyP production and facilitates industrial transfer, eventually contributing to the establishment of a circular P_i_ economy. Furthermore, a collection of clean deletion and overexpression strains was systematically compared under two cultivation conditions. This strain collection provides a platform for further analysis of phosphate metabolism in *S. cerevisiae* and corroborates many previously reported phenotypes in a distinct, uniform genetic background.

## 4. Methods

### 4.1 Molecular cloning

The CRISPR–Cas9 plasmid pV1382 (a gift from Gerald Fink; Addgene plasmid # 111436; http://n2t.net/addgene:111436; RRID: Addgene_111436; (Vyas et al., 2018), (Eerlings et al., 2024)), carrying a nourseothricin-resistance marker (NatR), was digested with BsmBI-v2 from New England Biolabs, Inc. (NEB) and dephosphorylated using recombinant shrimp alkaline phosphatase (rSAP; NEB). The linearized vector was separated by agarose-gel electrophoresis and purified using the Monarch DNA Gel Extraction Kit (NEB). Complementary single-guide RNA (sgRNA) oligonucleotides were phosphorylated with T4 polynucleotide kinase (NEB) and ligated into the digested vector using T4 DNA ligase (NEB). The ligation products were introduced by heat-shock transformation into NEB 5-alpha competent *E. coli* cells (DH5α; NEB). Transformants were selected on lysogeny broth (LB)-agar containing 100 mg/L ampicillin, and insertion of the correct sgRNA sequence was verified by Sanger sequencing.

To construct the repair templates, two homology arms of approximately 500 bp upstream and downstream of the coding sequence were amplified separately from CEN.PK113-7D genomic DNA using CloneAmp HiFi PCR Premix (Takara Bio, Shiga, Japan), gel purified and subsequently joined by overlap-extension PCR. The resulting repair template was gel-purified and reamplified to obtain sufficient DNA for transformation.

For chromosomal gene overexpression, integrative donor plasmids were constructed using pUC19 as the cloning backbone for propagation in *E. coli*. Each donor construct comprised approximately 500 bp upstream and downstream homology arms corresponding to sequences adjacent to the CRISPR–Cas9 cleavage site at the selected genomic integration locus. The expression cassette positioned between these homology arms consisted of approximately 600 bp promoter, the complete coding sequence of the gene to be overexpressed and approximately 250 bp terminator. All insert fragments were amplified from genomic DNA. The identities of the integration loci, promoters, terminators and overexpressed genes are provided in Supplementary Table S2.

The pUC19 backbone and individual insert fragments were assembled by Gibson assembly. Primers introduced 30 bp terminal overlaps homologous to the directly preceding and following DNA fragments, enabling ordered assembly of the complete donor plasmid. Assembly reactions were transformed into NEB 5-alpha competent *E. coli* cells (DH5α; New England Biolabs, Ipswich, MA, USA), and transformants were selected on lysogeny broth agar containing 100 mg/L ampicillin. Plasmid DNA was isolated from selected colonies using the Monarch Plasmid Miniprep Kit (New England Biolabs). Candidate constructs were initially screened for the expected plasmid size, and their complete sequences were subsequently verified by whole-plasmid sequencing (Eurofins Genomics Germany GmbH, Ebersberg, Germany). Only sequence-confirmed donor plasmids were used for subsequent strain construction.

### 4.2 *S. cerevisiae* strain construction

Markerless gene deletions and insertions were generated in *S. cerevisiae* CEN.PK113-7D using CRISPR– Cas9-mediated genome editing. All strains used in this study are listed in Supplementary Table S1. Genomic sequences of the target loci were obtained from the CEN.PK reference genome available through the *Saccharomyces* Genome Database (Engel et al., 2025; Nijkamp et al., 2012). sgRNAs targeting the respective coding sequences were designed using CRISPOR (Concordet and Haeussler, 2018) and complementary oligonucleotides containing BsmBI-compatible overhangs were designed for each selected guide. All primer sequences are provided in Supplementary Table S3.

Frozen competent yeast cells were prepared using a modified lithium acetate/single-stranded carrier DNA/polyethylene glycol (PEG) procedure (Gietz and Schiestl, 2007; Gietz et al., 1995). An overnight CEN.PK113-7D culture was used to inoculate 50 mL of 2× YPD medium supplemented with 80 mg/L adenine hemisulfate and cultivated at 30°C and 200 rpm to an OD₆₀₀ of 1.0–1.3. Cells were harvested at 3500 × g for 1 min, washed twice with sterile water and resuspended in 2 mL of frozen competent cell solution containing 5% (v/v) glycerol and 10% (v/v) dimethyl sulfoxide. Aliquots of 500 µL were frozen at approximately −1°C/min and stored at −80°C.

For each transformation, competent cells were thawed for 30 s at 42°C and separated into 100 µL aliquots, followed by centrifugation at 3500 × g. The supernatant was discarded, and the cell pellet was resuspended in 14 µL of DNA solution containing 600 ng of the pV1382-sgRNA plasmid and 1 µg of the corresponding repair template. Subsequently, 270 µL of transformation mixture containing 150 µL of 50% (w/v) PEG 3350, 45 µL of 1 M lithium acetate, and 75 µL of heat-denatured salmon-sperm carrier DNA at 2 mg/mL was added. Following vortex mixing, cells were heat-shocked at 42°C for 20 min, collected at 5000 × g for 1 min, resuspended in 1 mL of 5 mM CaCl₂ and incubated for 5 min. Cells were subsequently pelleted, and the supernatant was discarded. Cells were resuspended in 1 mL of yeast extract peptone dextrose (YPD) medium and recovered for 3.5 h at 30°C and 800 rpm before plating on YPD agar containing 50 mg/L nourseothricin (Jena Bioscience, Jena, DE).

Selected transformants were streaked to single colonies on nonselective YPD agar and subsequently cultivated overnight in liquid YPD to facilitate loss of the CRISPR plasmid. Plasmid curing was confirmed by the absence of growth on YPD agar containing 50 mg/L nourseothricin. Deletion of the target coding sequence was initially assessed by colony PCR using Phire Plant Direct PCR Master Mix (Thermo Fisher Scientific, Waltham, MA, USA) and a verification primer located outside the repair-template homology region. Amplicons of the expected size were verified by Sanger sequencing across the edited locus. Multiple independently isolated and sequence-confirmed deletion clones were retained and used as biological replicates in subsequent experiments.

Because repeated attempts to generate markerless *ARG82* and *KCS1* deletions using the NatR-selected CRISPR–Cas9 system did not yield verified deletion clones, the corresponding KanMX disruption cassettes were amplified from strains of the Yeast Knockout Collection (Giaever and Nislow, 2014) and transformed into CEN.PK113-7D. Transformants were selected on YPD agar containing 200 mg/L G418 (El Alami et al., 2003) and verified by colony PCR and Sanger sequencing.

For the integration of genes, the respective expression cassette was amplified from the corresponding plasmid and used as a repair template. A set of published target sites was used for integrations (Reider Apel et al., 2017).

### 4.3 Microscale cultivations following the overplus protocol

For precultures, the cells were inoculated from single colonies on YPD agar plates into 0.5 mL of synthetic defined (SD) medium in 96-well deep-well plates (System Duetz, Enzyscreen, Bennebrouek, NE) and incubated overnight at 30°C, 5 cm shaking diameter and 250 rpm. For biomass production, 20 µL of each overnight culture was transferred into 2.5 mL of SD medium in white 24 well clear-bottom plates (Enzyscreen, Bennebrouek, NE). Cultures were incubated for 24 h at 30°C, 5 cm shaking diameter, and 250 rpm. Culture volumes required for subsequent cultivation stages were calculated from green value (GV) signals using an empirically established calibration between GV and OD_600_. For P_i_ starvation, a volume corresponding to 7.5 OD_600_mL was collected by centrifugation at 5000 × g. Cells were washed once with 1 mL of ultra-pure water and resuspended in 1 mL of P_i_ starvation medium. The suspension was transferred to a 24-well plate containing 1.5 mL of starvation medium, resulting in a final cultivation volume of 2.5 mL per well. Cultures were incubated for 6 h at 30°C, 5 cm shaking diameter, and 200 rpm.

For P_i_ refeeding, feeding medium was prepared immediately before use by combining nine volumes of glucose–KH₂PO₄ solution with one volume of 200 mM MgCl₂ solution and equilibrating the mixture to 30°C. Following starvation, a culture volume corresponding to 6.5 OD_600_mL was collected at 5000 × g and washed once with 1 mL of ultra-pure water. Cells were resuspended in 0.3 mL of feeding medium and transferred to gray 96-well Growth Profiler plates. Refeeding cultures were incubated for 2.5 h at 30°C and 200 rpm.

At harvest, a volume of 250 µL was collected, cells were pelleted at 5000 × g, washed twice with 1 mL of ultra-pure water, and resuspended in 80 µL of ME buffer [25 mM MOPS, 2.5 mM EDTA]. The final biomass concentration was calculated from the GV, and the suspensions were stored frozen at −20°C until extraction (Christ and Blank, 2019).

### 4.4 Microscale cultivations following the P_i_-replete protocol

Precultures were inoculated in 0.5 mL of SD-medium in 96-well deep-well plates (System Duetz, Enzyscreen, Bennebrouek, NE) and incubated overnight at 30°C and 250 rpm. For biomass production, 20 µL of each overnight culture was transferred into 2.5 mL of SD medium (initial screening, Fig. 2B) or buffered Verduyn medium (P_i_ concentrations, Fig. 3C and D) in gray 24-well clear-bottom plates (Enzyscreen, Bennebrouek, NE). Cultures were incubated for 24 h at 30°C and 250 rpm.

At harvest, a volume of 1.2 mL was collected, cells were pelleted at 5000 × g, washed twice with 1 mL of ultra-pure water, and resuspended in 80 µL of ME buffer (25 mM MOPS, 2.5 mM EDTA). The final biomass concentration was estimated from the GV, and the suspensions were stored frozen at −20°C until extraction.

### 4.5 Cultivation in shake flasks

Shake flask cultivation under P_i_-replete and overplus conditions was done according to Christ and Blank (2019).

### 4.6 Analysis of growth rate and GFP expression

Growth and GFP fluorescence were monitored online using a BioLector microbioreactor system (Beckman Coulter GmbH, Aachen, Germany). Precultures were cultivated in 0.5 mL of minimal medium in 96-well deep-well plates (Enzyscreen, Bennebrouek, NE) at 30°C and 250 rpm. Main cultures were inoculated from the precultures to an initial OD₆₀₀ of 0.2 in 800 µL of buffered Verduyn medium in 48-well FlowerPlates (Beckman Coulter GmbH, Aachen, Germany). Cultivation was performed at 30°C and 1500 rpm with relative humidity maintained at 85%. Biomass formation was monitored as scattered light at 620 nm using a gain of 50. GFP fluorescence was measured at excitation and emission wavelengths of 488 nm and 520 nm, respectively, using a gain of 100.

Specific growth rates (µ) were determined from three biological replicates using QurvE (Wirth et al., 2023). Linear regression analysis of the natural logarithm of the scattered-light signal against cultivation time during exponential growth was performed and is reported in h⁻¹. For GFP evaluation, fluorescence values were normalized to the corresponding scattered-light signal to account for differences in biomass.

### 4.7 Fermentation in stirred tank reactors

#### 4.7.1 Precultures for bioreactor fermentations

*S. cerevisiae* strains were streaked from cryogenic stocks onto YPD agar plates and incubated at 30°C for 48–72 h. A single colony was used to inoculate 500 mL YPD medium containing 20 g/L glucose in a 1.8 L Fernbach flask. Pre-cultures were grown at 30°C and 100 rpm (100 mm throw) to an OD_600_ of 20, harvested by centrifugation at 4500 × *g* (CR22N, Eppendorf AG, Hamburg, Germany), and resuspended in 50 mL ultrapure water for bioreactor inoculation.

#### 4.7.2 Fermentation in DASBox

Initial strain characterization under fed-batch conditions was performed in an 8-fold-parallel DASbox Mini Bioreactor System equipped with a dual-head bioprocess autosampler for microbiological applications (Eppendorf AG, Hamburg, Germany). The system was controlled by DASware control v6.6.1. Each vessel had a total volume of 0.35 L and was operated with an initial working volume of 0.10 L. The bioreactors were equipped for online data acquisition, including a dissolved oxygen (DO) probe (VisiFerm DO ECS 120 H0, Hamilton, Bonaduz, Switzerland), a pH probe (EasyFerm Plus PHI K8 120, Hamilton, Bonaduz, Switzerland), and a Pt100 temperature sensor. Cultivations were carried out at 30°C and pH 5.0, controlled by adding 10% (w/w) NH_4_OH. DO was kept above 30% by automatically adjusting the stirring rate from 600 to 2000 rpm while airflow remained constant at 9 L/h. Off-gas was dried using a condenser; O_2_ and CO_2_ concentrations were monitored for the calculation of oxygen transfer rate, carbon transfer rate, and RQ (DASGIP GA4 Exhaust Analyzing Module, Eppendorf AG, Hamburg, Germany). The agitation shaft featured two six-blade Rushton turbines (⌀ 29 mm). Modified synthetic Verduyn medium contained (per L) 5 g (NH_4_)_2_SO_4_, 6.7 g KH_2_PO_4_, 4.0 g MgSO_4_ × 7 H_2_O, 20 mL Verduyn trace elements (TE) solution, and 2 mL Verduyn trace vitamins (TV) solution. The bioreactor was inoculated from a YPD preculture to an OD_600_ of 10-20, and exponential feeding was initiated immediately after inoculation. The feed solution contained (per L) 148 g glucose, 36.9 g KH_2_PO_4_, 33.4 g MgSO_4_ × 7 H_2_O, 10 mL TE, and 1 mL TV solution. During the fed-batch, the feed was pumped into the bioreactor via a precision peristaltic pump (DASGIP MP8 pump module, Eppendorf AG, Hamburg, Germany) equipped with Bioprene tubing (I.D. 0.5/W 1.05 mm). The feed rate was regulated via scripting in DASware control software using Equation 1, resulting in flow rates ranging from 0.8 to 15.9 mL/h.

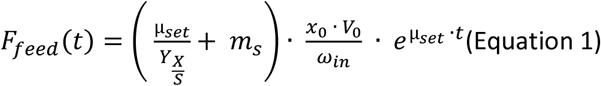

In Equation (1), *F_feed_* represents the feed rate in L/h, *x_0_* denotes the initial biomass concentration in g/L, ω_in_ indicates the glucose concentration in the feed in g/L, *V_0_* is the initial reactor volume, and *t* refers to the time in hours after the feeding begins. *Y_X/S_* is the maximum biomass yield on glucose without maintenance in g_CDW_/g_glucose_. *m_s_* is the maintenance coefficient in g_glucose_/g_CDW_/hour. The specific growth rate (µ_set_) was kept at 0.15 1/h. In total, 14.8 g of glucose (0.10 L feed) was fed in all fed-batch cultivations.

#### 4.7.3 Optical density measurement

Yeast growth was determined by measuring the optical density (OD_600_) at 600 nm with the Ultrospec 10-cell density meter (Amersham Biosciences, Amersham, UK). A correlation between OD_600_ and cell dry weight (CDW) was established.

#### 4.7.4 Cell dry weight determination

To determine the CDW, a 2 mL sample of the culture broth was centrifuged at 4°C and 15,000 × *g* for 2 min. The pellet was then resuspended in 1 mL of ultrapure water and centrifuged again under the same conditions, discarding the supernatant afterward. The pellet was suspended in another 1 mL of ultrapure water and transferred to a pre-weighed, pre-dried HPLC glass vial. This vial was dried at 65°C for 48 h, after which the CDW was calculated by weighing the vial.

#### 4.7.5 Metabolite quantification via HPLC

Quantification of the following metabolites: glucose, ethanol, glycerol, and acetate was performed using an Ultimate 3000 HPLC system (Thermo Scientific, Waltham, MA, USA). For this, an isocratic method with a mobile phase of 5 mM H_2_SO_4_ and a flow rate of 0.6 mL/min was used. A sample of 5 µL was injected onto a Metab-AAC column (ion exchange; dimensions: 300 × 7.8 mm; particle size: 10 µm; Isera GmbH, Düren, Germany) that was maintained at 40°C. The metabolites were detected by refractive index and UV detector (210 nm) and eluted in the following order: glucose at 9.3 min, glycerol at 13.5 min, acetate at 15.3 min, and ethanol at 21.8 min.

#### 4.7.6 Preparation of cells for polyP quantification

For lyophilization of yeast cells, a defined volume of culture broth corresponding to 6.8 mg CDW was centrifuged in a 2.0 mL pre-weighed reaction tube. The pellet was washed twice with ultrapure water, frozen at −80°C, and subsequently lyophilized at below 20 mbar for 24 h. The CDW was determined gravimetrically from the difference in tube weight before and after drying. For analytical polyP extraction, the dry cells were resuspended in 400 µL of ME buffer (25 mM MOPS, 2.5 mM EDTA) and frozen at −20°C.

### 4.8 Analytical polyP extraction

For polyP extraction, the samples were thawed, vortexed for 20 s, and briefly centrifuged at 1000 × g. Subsequently, 60 µL (microscale cultivations) or 300 µL (Bioreactor and shake flask) of saturated phenol was added, and the samples were vortexed for 20 s. Subsequently, samples were incubated at 45°C for 10 min, cooled on ice for 2 min, and briefly centrifuged at 1000 × g.

After addition of 200 µL (microscale cultivations) or 1 mL (bioreactor and shake flask) of chloroform, samples were vortexed for 20 s and briefly centrifuged at 1000 × g. Approximately 200 mg of high-vacuum silicone grease (Molykote, Dow Corning) was placed in the tube cap, and phase separation was achieved by centrifugation at 12,000 × g for 5 min. The upper aqueous phase containing polyP was transferred to a fresh tube, and the recovered extract volume was recorded. Extracts were stored at −20°C until quantification. For CEN.PK113-7D and most deletion strains, extracts were sequentially diluted to meet the criteria for quantification.

### 4.9 PolyP quantification

Quantification of polyP and P_i_ was done using a commercial kit (Phosfinity Quant, Aminoverse B.V., Nutz, NE) according to the Kit instructions (Christ and Blank, 2018).

### 4.10 Electron microscopy

Cells were grown in 50 mL of buffered Verduyn medium to an OD_600_ of 1 and harvested by centrifugation at 3,000 × g. Cells were fixed with 3% glutaraldehyde in 0.1 M Sørensen’s phosphate buffer, embedded in 1% low-melting agarose (Sigma, Steinheim, Germany), frozen in a high-pressure freezer (EM ICE, Leica, Wetzlar, Germany) and embedded into epon (Serva, Heidelberg, Germany) via freeze substitution. For this, samples were incubated in ethanol at −80°C for 50 h, brought slowly to room temperature, incubated in fresh ethanol 3 times for 10 min, 30 min in propylene oxide, 2 h in a 1:1 mixture of propylene oxide and epon, 2 h in pure epon and finally cured in pure epon for 2 h at 90°C.

Ultrathin sections (70-100 nm) were stained with 0.5% uranyl acetate and 1% lead citrate (both EMS, Munich, Germany) to enhance contrast. Samples were examined using a Hitachi HT7800 transmission electron microscope (Hitachi, Tokyo, Japan) operating at an acceleration voltage of 100 kV.

For EDX analysis 1 µm-thick sections were placed on silicon wafers, stained with 4% ytterbium acetate (Sigma-Aldrich, St. Louis, USA) and 1% lead citrate and coated with 10 nm carbon. Elemental analysis was performed with a scanning electron microscope Quattro S (Thermo Fisher, Eindhoven, Netherlands) at 4 kV accelerating voltage in a high-vacuum environment.

### 4.11 Media compositions

TE solution contained (per L): 1.50 g Na_2_EDTA × 2 H_2_O, 0.45 g ZnSO_4_ × 7 H_2_O, 0.10 g MnCl_2_ × 2 H_2_O, 0.03 g CoCl_2_ × 6 H_2_O, 0.03 g CuSO_4_ × 5 H_2_O, 0.04 g Na_2_MoO_4_ × 2 H_2_O, 0.45 g CaCl_2_ × 2 H_2_O, 0.30 g FeSO_4_ × 7 H_2_O, 0.10 g H_3_BO_3_, and 0.01 g KI.

TV solution contained (per L): 0.05 g D-biotin, 1.00 g Ca-D-pantothenate, 1.00 g nicotinic acid, 25.00 g myo-inositol, 1.00 g thiamine HCl, 1.00 g pyridoxine HCl, 0.20 g *p*-aminobenzoic acid.

Verduyn medium contained (per L): 20 g glucose, 5 g (NH_4_)_2_SO_4_, 3 g KH_2_PO_4_ (for the P_i_-containing version), 0.5 g MgSO_4_ × 7H_2_O, 20.4 g potassium hydrogen phthalate (for the buffered version), 1 mL TV, 10 mL TE, pH adjusted to 5.2 with 1 M KOH, and the medium was sterile-filtered.

SD medium contained (per L): 6.70 g yeast nitrogen base without amino acids (Sigma-Aldrich), 20.00 g glucose. The pH was adjusted to 5.4, and the medium was sterile-filtered.

Starvation medium contained (per L): 23.96 g glucose, 1.32 g (NH_4_)_2_SO_4_, 0.99 g KCl, 0.58 g CaCl_2_, 2.02 g Na_2_Succinate, 10 mL TE solution, 1 mL TV solution, pH adjusted to 5 with HCl, and the medium was sterile-filtered.

Feeding medium contained (per L): 45 g glucose, 8.15 g KH_2_PO_4_, 4.06 g MgCl_2_×6H_2_O, pH adjusted to 6 with HCl and KOH, and the medium was sterile-filtered.

### 4.12 Statistics

Statistical analyses and graphical representations were performed using GraphPad Prism (Version 8.0.2). Data are presented as individual biological replicates with the mean ± SD, unless stated otherwise. For the strain-screening experiments, differences between strains were analyzed by ordinary one-way ANOVA followed by Dunnett’s multiple-comparisons test against the corresponding wild-type control. Experiments examining the effects of strain and P_i_ starvation treatment were analyzed by ordinary two-way ANOVA, followed by Tukey’s multiple-comparisons test. Model assumptions were evaluated by inspection of residual and quantile–quantile plots.

## Supporting information

supplementary graphs

Strains, Plasmids, Primers

## 5. CRediT author-contribution statement

Makarius Baier: Conceptualization, Data curation, Formal analysis, Investigation, Methodology, Resources, Supervision, Validation, Visualization, Writing – original draft, Writing – review & editing; Philipp Demling: Conceptualization, Funding acquisition, Investigation, Methodology, Project administration, Supervision, Writing – review & editing; Lars M. Blank: Conceptualization, Funding acquisition, Project administration, Resources, Supervision, Writing – review & editing; Tobias Karmainski: Conceptualization, Data curation, Formal analysis, Investigation, Methodology, Validation, Visualization, Writing – original draft, Writing – review & editing; Eva Miriam Buhl: Data curation, Investigation, Methodology, Resources, Writing – review & editing; Alexander Deitert: Conceptualization, Investigation, Methodology, Writing – review & editing; Roy Eerlings: Methodology, Resources, Writing – review & editing; Evelyn Deis: Investigation; Frederik Völker: Methodology, Writing – review & editing; Lennart Brans: Investigation, Methodology

## 7. Acknowledgments

The authors thank their colleagues at the Institute of Applied Microbiology (iAMB) for their good spirits and continued support. The authors thank Eppendorf for providing the DasBox with Dual-Head bioprocess autosampler and the VH Berlin (research institute for bakers’ yeast) for providing the strain *S. cerevisiae* VH2.200.

## 9. AI-use declaration

During the preparation of this work, the authors used ChatGPT (OpenAI) to support language refinement. After using this tool, the authors reviewed, verified, and edited the content as needed and take full responsibility for the content of the article.

## 10. Data-availability statement

Data will be made available on request.

