## supplementary graphs for "Metabolic engineering of *Saccharomyces cerevisiae* for improved polyphosphate production"

**Supplementary material "Metabolic engineering of *S. cerevisiae* for improved polyphosphate production"**

### S1. *PHO80* deletion moderately reduces growth in buffered Verduyn medium

To assess the effect of *PHO80* deletion on growth under the conditions used for orthophosphate ($P_i$)-replete polyP accumulation, CEN.PK113-7D wild type (WT) and *pho80Δ* were cultivated in buffered Verduyn medium and biomass formation was monitored online by backscatter (Fig. S1). The *pho80Δ* strain exhibited a delayed increase in biomass and reached a lower final backscatter signal than the WT. Analysis of the exponential-growth phase yielded maximum specific growth rates of 0.40 ±0.03 $h^{-1}$, and 0.31 ±0.02 $h^{-1}$, respectively. Thus, *PHO80* deletion imposed a moderate growth penalty without preventing rapid growth under $P_i$-replete conditions.

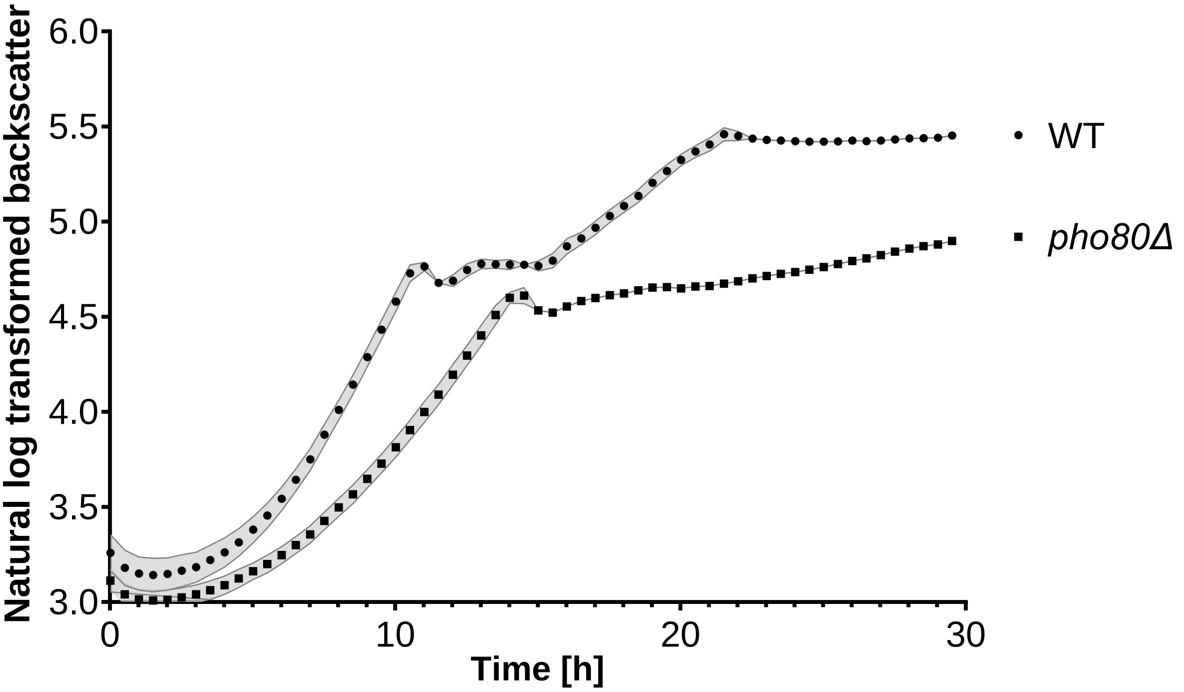


*Fig. S1: Growth of S. cerevisiae CEN.PK113-7D wild type (WT) and pho80Δ in buffered Verduyn medium. Biomass formation was monitored online by backscatter during 30 h of cultivation. Curves show mean natural-log-transformed backscatter values, and shaded areas represent mean ± SD from three independent biological replicates (n = 3).*

#### S2. Residual $P_i$ after cultivation at different initial $P_i$ concentrations

To determine the proportion of supplied $P_i$ remaining after cultivation, residual $P_i$ was quantified in the culture supernatants of *pho80Δ* grown in buffered Verduyn medium containing different initial $P_i$ concentrations (Fig. S2). At initial concentrations up to 7.9 mM, residual $P_i$ was below the analytical quantification limit, indicating near-complete consumption of the supplied $P_i$. Residual $P_i$ became detectable at 8.8 mM and increased strongly when 22 mM $P_i$ was supplied. These results complement the cellular polyP measurements in Fig. 3C and demonstrate that the plateau in polyP accumulation at high $P_i$ supply is accompanied by incomplete uptake of the supplied $P_i$.

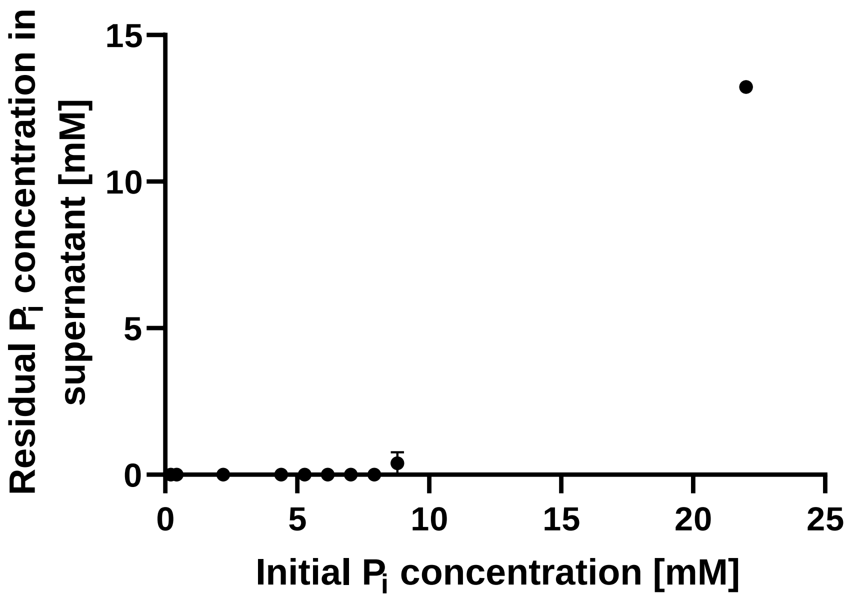


***Fig. S2: Residual phosphate ($P_i$) after cultivation of CEN.PK113-7D pho80Δ at different initial $P_i$ concentrations***. *Cells were cultivated in buffered Verduyn medium containing the indicated initial $P_i$ concentrations, and residual $P_i$ in the supernatant was quantified after 24 h. Data are presented as mean ± SD from three independent biological replicates. Values below the analytical quantification limit were plotted as zero.*

**S3. Individual biological replicates confirm constitutive PHO5 promoter activity in *pho80Δ***

To evaluate the reproducibility of the *PHO5* promoter response presented in Fig. 4, the trajectories from the two independent biological experiments are shown separately in Fig. S3. In the WT, reporter activation depended strongly on the initial $P_i$ concentration and occurred after a concentration-dependent delay. Despite variation in the timing and magnitude of the response between the two experiments, the reporter was preferentially induced when $P_i$ availability became limiting. By contrast, biomass-normalized GFP fluorescence increased in *pho80Δ* across the complete range of initial $P_i$ concentrations. The individual trajectories therefore support constitutive activation of the *PHO5* promoter following *PHO80* deletion.

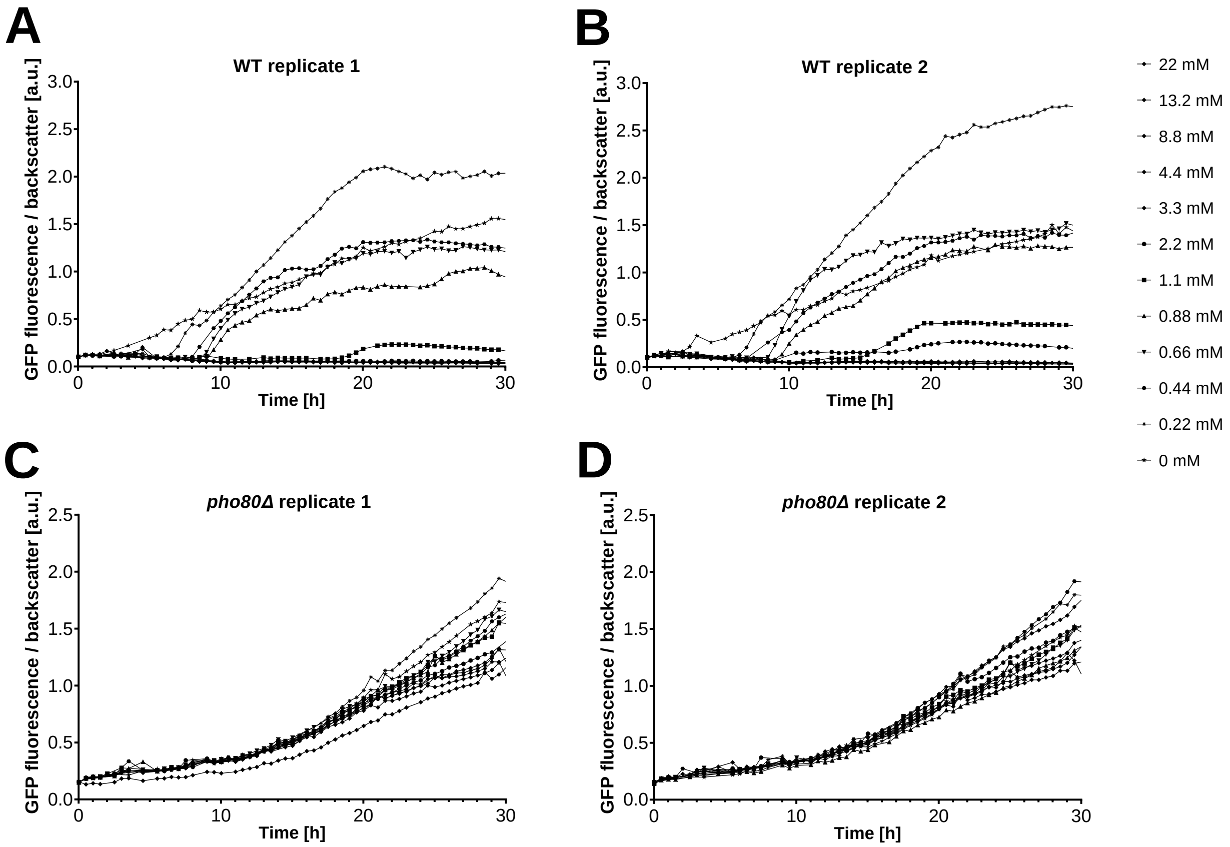


***Fig. S3: Individual biological replicates of the PHO5 promoter reporter experiment presented in Fig. 4**. CEN.PK113-7D wild type (WT) (**A** and **B**) and pho80Δ (**C** and **D**) carrying an integrated ars416d::$P_{PHO5}$–sfGFP reporter cassette were cultivated in buffered Verduyn medium containing initial $P_i$ concentrations between 0 and 22 mM. GFP fluorescence was normalized to the corresponding backscatter signal. Each panel represents one independent biological experiment, and each trajectory represents the indicated initial $P_i$ concentration.*

**S4. Functional validation of constitutive GFP expression cassettes**

The functionality of the chromosomally integrated expression cassettes was evaluated using sfGFP controlled by the constitutive *TEF1* or *TDH3* promoter (Fig. S4). Both engineered strains produced substantially higher GFP fluorescence than the non-fluorescent WT control, confirming functional expression from both integrated cassettes. Their backscatter profiles remained comparable to that of the parental strain, indicating that cassette integration and constitutive sfGFP expression did not cause an evident growth defect under the tested conditions.

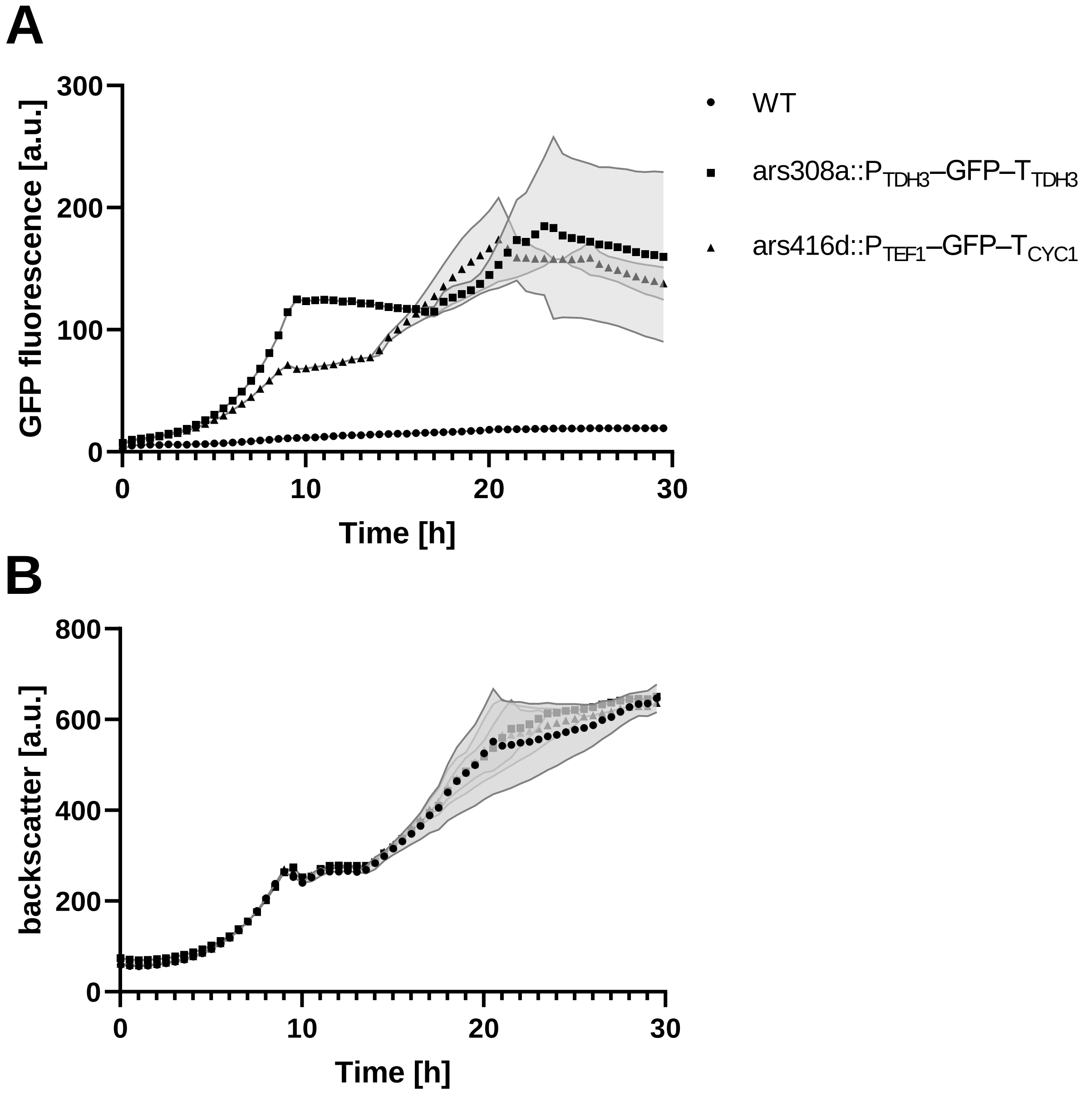


***Fig. S4: Functional validation of constitutive sfGFP expression cassettes in S. cerevisiae CEN.PK113-7D**. GFP fluorescence (**A**) and backscatter (**B**) were monitored in the wild type (WT) and strains carrying ars416d::P$_{TEF1}$-sfGFP-T$_{CYC1}$ or ars308a::P$_{TDH3}$-sfGFP-T$_{TDH3}$ expression cassettes. Curves show mean values, and shaded areas indicate SD from three independent biological replicates. The WT served as an autofluorescence control.*
